# Liquid-like VASP condensates drive actin polymerization and dynamic bundling

**DOI:** 10.1101/2022.05.09.491236

**Authors:** Kristin Graham, Aravind Chandrasekaran, Liping Wang, Aly Ladak, Eileen M. Lafer, Padmini Rangamani, Jeanne C. Stachowiak

## Abstract

The organization of actin filaments into bundles is required for cellular processes such as motility, morphogenesis, and cell division. Filament bundling is controlled by a network of actin binding proteins. Recently, several proteins that comprise this network have been found to undergo liquid-liquid phase separation. How might liquid-like condensates contribute to filament bundling? Here, we show that the processive actin polymerase and bundling protein, VASP, forms liquid-like droplets under physiological conditions. As actin polymerizes within VASP droplets, elongating filaments partition to the edges of the droplet to minimize filament curvature, forming an actin-rich ring within the droplet. The rigidity of this ring is balanced by the droplet’s surface tension, as predicted by a continuum-scale computational model. However, as actin polymerizes and the ring grows thicker, its rigidity increases and eventually overcomes the surface tension of the droplet, deforming into a linear bundle. The resulting bundles contain long, parallel actin filaments that grow from their tips. Significantly, the fluid nature of the droplets is critical for bundling, as more solid droplets resist deformation, preventing filaments from rearranging to form bundles. Once the parallel arrangement of filaments is created within a VASP droplet, it propagates through the addition of new actin monomers to achieve a length that is many times greater than the initial droplet. This droplet-based mechanism of bundling may be relevant to the assembly of cellular architectures rich in parallel actin filaments, such as filopodia, stress fibers, and focal adhesions.

## INTRODUCTION

VASP is an actin filament nucleator and polymerase that catalyzes the processive elongation of actin filaments at the leading edge of cells, in focal adhesions, and at filopodial tips ^1^. VASP plays a significant role in cell motility, adhesion, and sensing. It is part of the Ena/VASP family of proteins that share a conserved modular structure. In particular, VASP is a homo-tetramer, where each monomer consists of an N-terminal EVH1 domain, a central proline-rich region, and a C-terminal EVH2 domain (Figure 1A) ^2^. The EVH2 domain consists of G-actin and F-actin binding domains, which together confer the polymerase activity of VASP, as well as a coiled-coil domain that mediates tetramerization ^3, 4^.

**Figure 1:**
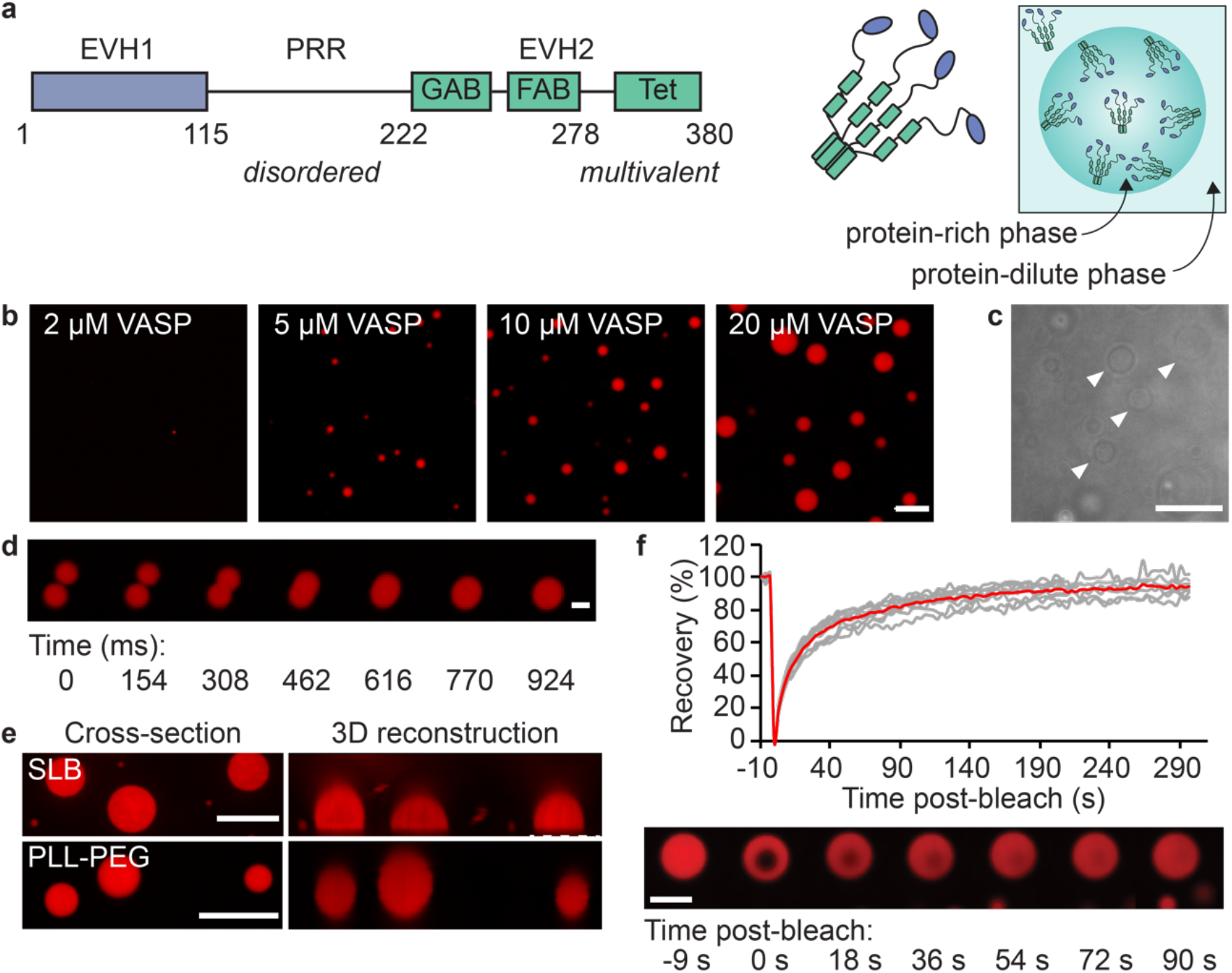
VASP forms liquid-like droplets in vitro. (a) *Left:* Domain organization of VASP. *Right:* Schematic of VASP tetramer and droplet formation. (b) Droplets formed by VASP (labeled with AlexaFluor-647) at increasing protein concentrations. The buffer used in these experiments, hereafter referred to as “droplet buffer”, contained 50 mM Tris pH 7.4, 150 mM NaCl, 0.5 mM EDTA/EGTA, 5 mM TCEP, and 3% (w/v) PEG8K. Scale bar 5 μm (c) Brightfield images of unlabeled 30 μM VASP droplets (arrows). Scale bar 10 μm. (d) Fusion and re-rounding of two VASP droplets. Scale bar 2 μm. © 20 μM VASP droplets exhibit wetting of glass coverslips passivated with supported lipid bilayers (SLB), but not PLL-PEG passivated coverslips. Scale bar 10 μm. (f) FRAP of 15 μM VASP droplets. *Top:* Recovery profile after photobleaching a 1.8 μm ROI within VASP droplets. The average recovery is 93.9 ± 4.89% (n=9 droplets). Red line indicates average, gray lines represent each independent droplet. *Bottom:* Image series of fluorescence recovery of VASP droplets. Scale bar 5 μm.

The capacity of the VASP tetramer to form multivalent networks with other actin-binding proteins suggests that it could provide a scaffold for assembly of protein networks. Indeed, VASP has been found to form dynamic protein assemblies both *in vivo* and *in vitro*. The proteins IRSp53 and lamellipodin have both been shown to recruit VASP to the plasma membrane, forming dynamic protein-rich clusters ^5, 6^. These clusters can function to localize actin polymerization and, in some cases, precede filopodia formation. VASP has also been found to cluster on actin filaments with lamellipodin ^7^. These assemblies of VASP are thought to be critical for the processive elongation of actin filaments, as clustering of VASP results in more robust actin polymerization ^8, 9^. Collectively, these studies suggest that protein assemblies rich in VASP help to organize the cytoskeleton in multiple cellular contexts.

Increasing evidence suggests that many cellular proteins are organized into liquid-like condensates through the process of liquid-liquid phase separation (LLPS) ^10–12^. Assembly of condensates is primarily mediated by weak, multivalent interactions between proteins with substantial disorder or conformational flexibility. These assemblies often exhibit liquid-like behavior, such as fusion, wetting, and high degrees of molecular rearrangement ^12^.

Recent work has shown that some cytoskeletal effectors are able to assemble into condensed phases mediated by LLPS, and these condensates are capable of driving polymerization of cytoskeletal filaments ^11, 13–15^. What role could protein droplets play in organizing cytoskeletal filaments into higher order assemblies? Here we show that VASP undergoes LLPS to form liquid-like droplets. These droplets strongly localize actin polymerization. To minimize curvature, actin filaments partition to the edges of the droplets, forming a growing ring. When the rigidity of the ring exceeds the mechanical constraints of the droplet surface tension, a linear actin bundle is formed through a series of symmetry breaking events. Guided by a continuum-scale computational model, this work describes a physical mechanism by which the unique, liquid-like properties of protein droplets can bundle cytoskeletal filaments and perpetuate their growth over long physiologically relevant length scales.

## RESULTS

### The tetrameric actin polymerase, VASP, forms liquid-like droplets

As a homo-tetramer, VASP has an inherent potential for multivalent interaction. Additionally, about 60% of VASP is predicted to be intrinsically disordered, Figure 1A, Supplemental Figure S1 ^13, 16, 17^. VASP’s intrinsically disordered region has a high proline content, which may facilitate interactions with other actin-interacting proteins ^18^. These features suggest that VASP may form extended multi-valent networks. To test this idea, we combined VASP (2-20 μM monomer concentration, ~10% labeled with Alexa-Fluor 647) with the crowding agent polyethylene glycol (PEG 8k, final concentration 3% (w/v)). Here PEG is included to mimic the crowded cytoplasm, a common practice in the LLPS field ^19–22^. Upon imaging with fluorescence confocal microscopy, the presence of micrometer-scale, spherical, protein-rich droplets were observed within 10 minutes of PEG addition, Figure 1B. The number and size of these droplets increased with the concentration of VASP at constant temperature, as expected for LLPS ^23, 24^. Unlabeled VASP formed similar droplets to those formed from labeled protein (Figure 1C). VASP droplets fused and re-rounded, within about one second following contact, indicating their fluid-like nature, Figure 1D, Supplemental Video S1. They wetted glass coverslips that were coated with hydrophilic lipid bilayers, becoming dome-like, providing further evidence of their fluidity, Figure 1E, top. In contrast, substrates coated with polyethylene glycol were repulsive to droplets, such that the droplets remained spherical, Figure 1E, bottom. To probe the rate of molecular exchange within droplets, we measured fluorescence recovery after photobleaching (FRAP). A region of about 2 μm in diameter within VASP droplets recovered nearly completely (fractional recovery 94 ± 5%) in less than two minutes (t1/2 18 ± 3 sec) (Figure 1F, Supplemental Video S2), indicating that molecular exchange was rapid within VASP droplets. To evaluate the molecular determinants of phase separation by VASP, we created a series of mutants and tested their ability to form droplets (Supplementary Figure S2). These studies revealed that the tetramerization domain of VASP is essential for phase separation, and that contacts between the tetramers, required for a long-range network, likely rely on weak, electrostatic interactions among multiple domains within VASP (Supplementary Figure 2).

### Polymerization of actin inside VASP droplets progressively transforms droplet shape

We next investigated the role of VASP droplets in actin polymerization and filament organization. We added G-actin monomers labeled with Atto-488 to protein droplets composed of VASP, labeled with AlexaFluor-647. We examined increasing ratios of actin to VASP including: 1:20 (1 μM actin, 20 μM VASP), 1:10 (2 μM actin, 20 μM VASP), and 1:7.5 (2 μM actin, 15 μM VASP). Under each of these conditions, actin localized strongly to VASP droplets, resulting in dramatic changes in droplet morphology, with many droplets taking on elongated shapes (Figure 2a). Specifically, about 10-15 minutes after addition of actin, we observed several distinct droplet morphologies, including: (i) a homogeneous distribution of actin within spherical droplets, (ii) peripheral accumulation of actin at the inner surfaces of spherical droplets, (iii) deformation of droplets into ellipsoid shapes, and (iv) rod-like, actin-filled droplets, Figure 2b. Notably, staining with phalloidin, which specifically binds filamentous actin, resulted in similar actin morphologies, indicating that these arrangements likely arise from polymerization of filaments, Figure 2c, *left*. Furthermore, in the presence of the polymerization inhibitor, latrunculin A, all droplets remained spherical in shape with uniform G-actin partitioning, suggesting that actin polymerization was responsible for the observed changes in droplet morphology, Figure 2c, *right*. Interestingly, the distribution of these morphologies changed as the actin to VASP ratio increased, Figure 2d. Specifically, for an actin to VASP ratio of 1:20, most VASP droplets were spherical with homogeneous or peripheral actin distributions. As this ratio increased to 1:10, elliptical and rod-like morphologies emerged, with elliptical droplets making up the majority. Finally, for a ratio of 1:7.5, most droplets had elongated shapes, with more than half displaying rod-like morphologies. Time-lapse imaging of VASP droplets after addition of actin revealed that actin rearranged within droplets from an initially homogeneous distribution to a peripheral distribution (Figure 2e and Supplemental Video S3) and that droplets with a peripheral distribution of actin frequently transformed into ellipsoid and rod-like morphologies, Figure 2f and Supplemental Video S4. Collectively these data suggest that actin polymerization within VASP droplets drives peripheral accumulation of actin, followed by a progressive increase in droplet aspect ratio, which leads to actin-filled droplets with elliptical and rod-like shapes. To better understand the impact of actin polymerization on droplet morphology, we sought to examine each of these transitions in greater detail, beginning with the peripheral accumulation of actin at the inner surfaces of VASP droplets.

**Figure 2:**
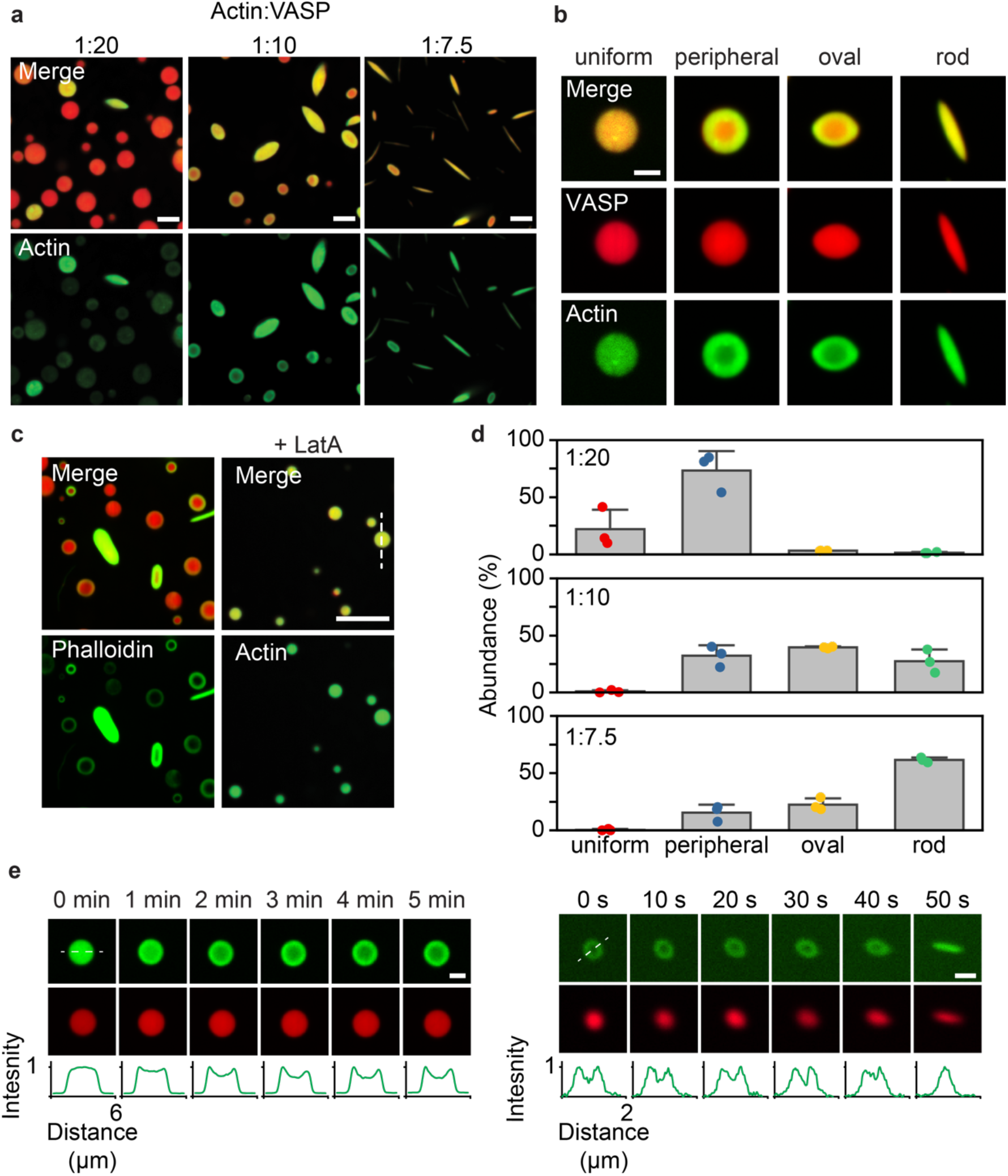
Actin polymerization within VASP droplets drive droplet deformation through discrete intermediates. (a) Increasing actin to VASP ratio results in increasingly elongated structures. Scale bar 5 μm. (b) Representative images of the different organizations of actin in VASP droplets. Scale bar 2 μm. (c) Phalloidin-iFluor-488 staining of VASP droplets to which unlabeled G-actin monomers were added. Pretreatment of VASP droplets with 5 μM LatrunculinA inhibits actin polymerization and results in the uniform partitioning of actin to VASP droplets. Scale bar 10 μm. (d) Quantification of the percent abundance of shapes observed across increasing actin to VASP ratios tested in A. Error bars represent standard deviation across n = 3 biologically independent experiments, with at least 600 droplets analyzed per replicate. (c) Time-lapse of peripheral actin accumulation in VASP droplets. Scale bar 2 μm. Line profile of actin intensity over time. Actin intensity is normalized to the max intensity in each frame. (f) Time-lapse of deformation of droplets with peripheral actin distribution to form rod-like droplets. Scale bar 1 μm. Line profile of actin intensity over time. Actin intensity is normalized to the max intensity in each frame.

### Actin polymerizes to form shells and rings inside VASP droplets

Shortly after the addition of actin, we observed individual actin filaments within VASP droplets, Figure 3a. As filaments grew within the droplets, they began to partition to the inner surfaces of the droplet. This peripheral distribution of actin appeared similar whether actin was labeled by inclusion of fluorescent-tagged monomers (Atto-488), or by phalloidin, Figure 3b, indicating that peripheral actin was filamentous. Strikingly, the peripheral accumulation of actin filaments resulted in assembly of three-dimensional actin shells and two-dimensional actin rings within the droplets, Figure 3c. Both structures appeared ring-like in individual confocal images but were distinguishable in three-dimensional reconstructions. In some cases, actin rings appeared to deform VASP droplets into two-dimensional discs. This observation suggests that as filaments accumulated at the droplet periphery and formed rings, their collective rigidity was sufficient to deform the droplet, Figure 3c, bottom. Among structures with peripheral actin, the distribution of shells, rings, and discs shifted as the actin to VASP ratio increased, from predominately shells at 1:20, to rings at 1:10, and to discs at 1:7.5. These data suggest that shells transformed into rings and then into discs as actin concentration increased, Figure 3d.

**Figure 3:**
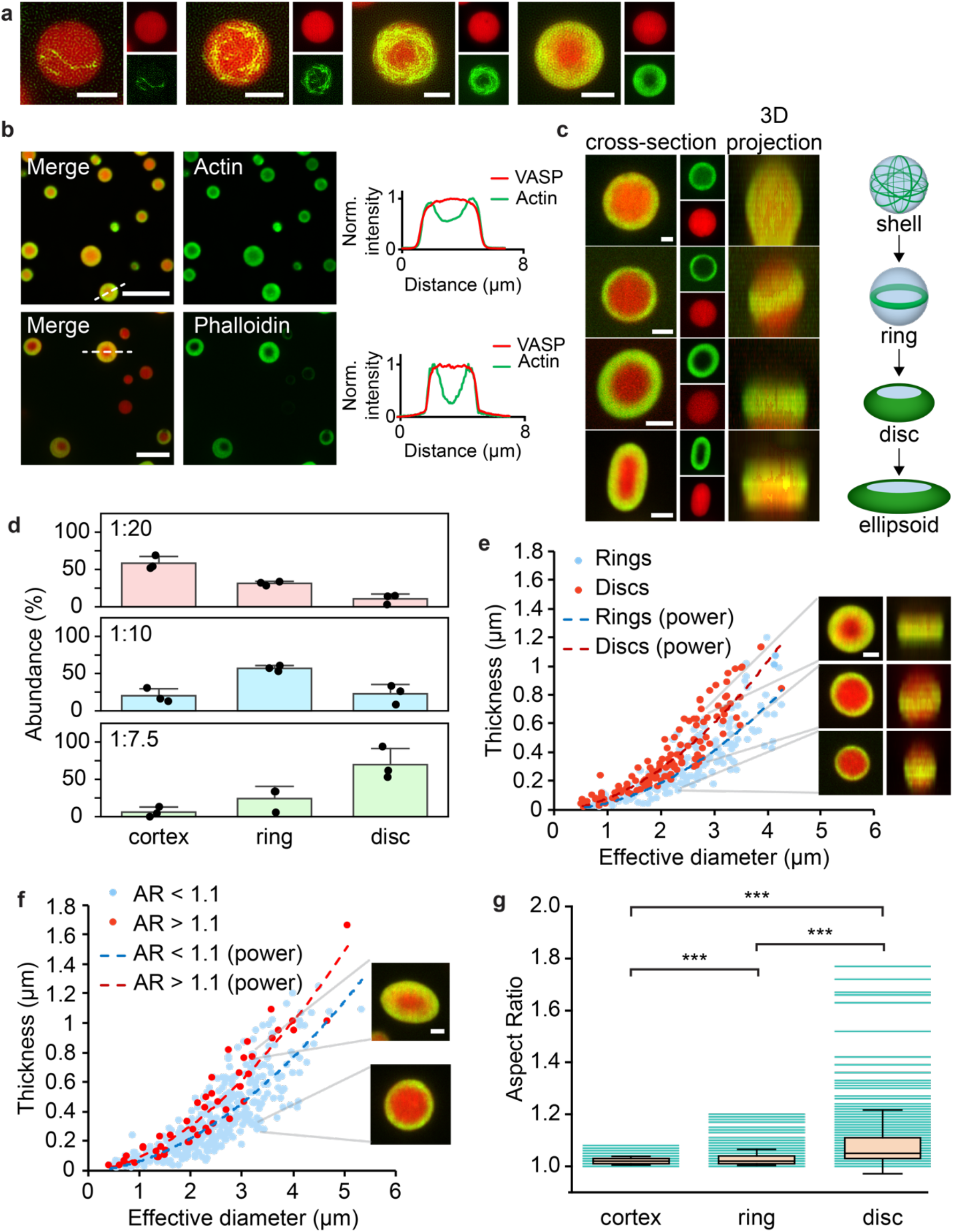
Peripheral accumulation of actin drives assembly of actin shells, rings, and discs. (a) Representative images of VASP (red) droplets containing an increasing amount of filamentous actin, as stained with phalloidin (green). Scale bar 2 μm. (b) *Top:* Addition of 1 μM actin to 20 μM VASP droplets results in the formation of a majority of peripheral actin within the droplet. Line profile of actin and VASP intensities inside the droplet. Scale bar 10 μm. *Bottom:* Phalloidin staining of peripheral actin within VASP droplets. Line profile of phalloidin-actin and VASP intensities inside the droplet. Scale bar 5 μm. Intensity is normalized to the max intensity of each channel. (c) Representative images of VASP droplets containing peripheral actin distribution. Peripheral actin distribution, as observed with phalloidin staining, can be further classified into shells, rings, or discs, as observed by 3D reconstructions. Scale bar 2 μm. (d) Quantification of the abundance of shells, rings, and discs as a function of actin to VASP ratio. Data are mean + standard deviation across n = 3 biologically independent experiments, with at least 170 droplets analyzed per condition©(e) Quantification of ring thickness as a function of droplet diameter, for disc and ring distributions of actin. Data is for droplets with aspect ratios < 1.1. Trendline represents the best fit to a power law of the form y = ax^b^. For rings, a = 0.0476 and b = 1.9682, R^2^ = 0.7278 (n=173 droplets). For discs, a = 0.0797, b = 1.8456, R^2^ = 0.8429 (n=131 droplets). Data from 3 biologically independent experiments. (f) Quantification of ring thickness as a function of droplet diameter, sorted by droplet aspect ratio. An effective diameter was calculated to allow comparison of droplet size between high and low aspect ratio structures (see methods). Trendline represents the best fit to a power law of the form y = ax^b^. For droplets with AR < 1.1, a = 0.0629 and b = 1.8037, R^2^ = 0.7111 (n=386 droplets). For droplets with AR > 1.1, a = 0.0917, b = 1.7315, R^2^ = 0.8979 (n=47 droplets). Data from 3 biologically independent experiments.(g) Distribution of droplet aspect ratios for droplets with shell, ring, or disc distributions of actin. Box represents interquartile range with whiskers representing standard deviation and median as a bisecting line. Each teal background line represents a single droplet. For shells, n = 167 droplets; for rings, n = 273 droplets; for discs, n = 227 droplets. Data collected from 3 biologically independent experiments. Brackets indicate data that was tested for significance using an unpaired, two-tailed t-test, with asterisks indicating p < 0.001.

Interestingly, the thickness of actin rings within spherical droplets increased with increasing droplet diameter (Figure 3e, Supplemental Figure S3). Further, for a given droplet diameter, droplets that had deformed into either discs (Figure 3e) or ellipsoids (Figure 3f, aspect ratio above 1.1) contained thicker rings in comparison to spherical droplets. Further, droplets that flattened into discs tended to have higher aspect ratios than spherical droplets (Figure 3g), suggesting that droplet elongation was coupled to the reduction in dimensionality associated with flattening. Overall, these data suggest that as the thickness of actin rings increased, the droplets began to deform. Specifically, as actin rings became thicker and more rigid, the energy required to confine them within droplets eventually exceeded the surface energy of the droplet, resulting in deformation. To better understand these mechanisms, we constructed a simple mechanical model.

### A continuum-scale computational model explains the mechanics of droplet deformation by actin

Filamentous actin has been previously observed to accumulate and form rings at the inner surfaces of soft, spherical containers such as membrane vesicles and aqueous/oil emulsions ^25, 26^. In these cases, partitioning of actin to the inner surfaces of the containers was observed whenever the diameter of the container was less than the persistence length of actin filaments, 10-20 μm. This criterion is met by the VASP droplets, which ranged in diameter from 1-7 μm, Supplemental Figure S4.

In the case of a growing filament within a deformable container, the relative stiffness of the filament determines whether it deforms the surface of the container or bends to fit within it ^27^. For filaments that remain within the container, the lowest energy state for the bent filament is reached when the filament partitions to the edge of the droplet, where it experiences the least curvature. This partitioning should lead to a shell of curved filaments at the inner surface of the droplet, as observed in Figure 3. The transition from a threedimensional shell to a two-dimensional ring (Figure 2c) has been previously observed in the presence of either macromolecular depletants or actin bundling proteins ^26^. Similarly, our observation of actin rings can be explained by both the crowded nature of the droplet interior and the ability of VASP to bundle actin filaments.

The total energy of the ring, including contributions from thermal fluctuations and filament bending, is given by Equation 1.

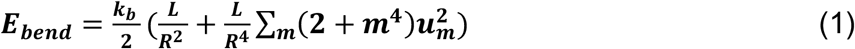

Here, *k_b_* is the bending rigidity of the ring, which consists of actin filaments crosslinked by VASP. 1/*R* is the filament curvature, where *R* is the droplet radius, and *L* is the filament length. The second term in Equation 1 represents the contributions from thermal fluctuations of the droplet-filament interface, where *m* is the Fourier mode of the fluctuations, and *u_m_* is the Fourier amplitude, given by equipartition theorem as, 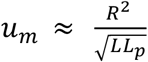. *L_p_* is the persistence length of the filaments. We assume that the thickness of the ring in a spherical droplet is proportional to the Fourier amplitude *u_m_* of the bundle. Hence, the droplets with larger ring thicknesses cannot remain spherical and will deform. See Supplementary materials for the full derivation ^25, 28^. This analysis, which builds on the work of Limozin et al ^25^, predicts that the ring thickness will increase in proportion to D^1.5^. This prediction is consistent with our experimental data, in which ring thickness scales as D^1.8^ (Figure 3f, droplets AR < 1.1). Consistent with the data and model predictions, studies of filament rings encapsulated within other types of spherical containers have also shown that the exponent ranges between 1.5 and 2. ^25, 28 29^

As noted above, actin polymerization was confined to VASP droplets and did not occur in the surrounding solution, Figure 2. This confinement implies that the wetting energy of actin in the VASP-rich phase, *γ_AVR_*, is significantly lower than the wetting energy of actin in the VASP-dilute phase, *γ_AVD_*. There is an additional energy component to consider, the interaction of VASP-rich droplet with the surroundings, *γ_I_*, which determines the shape of the droplet interface, effectively giving a surface energy for a droplet of surface area *A_I_*, as *E_I_* = *γ_I_A_I_*. As long as the ring bending energy, *E_bend_*, is less than *E_I_*, the droplet will retain its spherical shape. However, when *E_bend_* exceeds *E_I_*, the shape of the droplet must change to accommodate the excess bending energy of the actin ring compared to *E_I_*.

We modeled the interaction energies between the actin filaments and VASP droplets using a numerical energy minimization scheme (see Supplemental Figure S5 for details). For simplicity, we assume that the droplet is a 2D circle and the filament ring is represented by a curve with finite thickness (Figure 4a-b). Additionally, we assume that the ring thickness is uniform along the entire length of the ring, which is consistent with our images in Figure 2, thereby approximating the ring as a 1D curve. We establish the ring confinement within the droplet by tuning the values of *γ_AVR_* and *γ_AVD_*, Figure 4d, and Supplemental Figure S6). As we increased the ring thickness (Figure 4e bottom), for a fixed droplet radius and *γ_I_*, the shape of the droplet changed from a circle to an ellipse, Figure 4f. This behavior held for a range of droplet diameters, Figure 4g. To determine whether these relationships held for different values of *γ_I_*, we varied *γ_I_* for droplet diameters ranging from 0.5 – 4 μm. For each combination of droplet diameter and *γ_I_*, we plotted the aspect ratio of the resulting droplet as a function of increasing ring thickness, Figure 4g (additional *γ_I_* values in Supplemental Figure S7). Additionally, to identify the threshold of droplet deformation, we plotted the ring thickness above which the circular droplet began to deform (aspect ratio greater than 1.1) as a function of droplet diameter for a range of *γ_I_*. Here we found that the threshold ring thickness increased non-linearly with increasing droplet diameter (Figure 4h), in good agreement with the experimental data in Figure 3e. The resulting curve can be thought of as a phase boundary. For each value of droplet diameter, it tells us the critical ring thickness, above which a spherical droplet will begin to elongate into an ellipsoid. As droplet radius increases, surface energy increases, such that thicker rings are required to deform droplets. Additionally, for a given diameter, droplets with high surface energy can accommodate thicker rings without deforming. Overall, the model-data comparison suggests that deformation of droplets is governed by a competition between actin bending energy and droplet surface energy.

**Figure 4.**
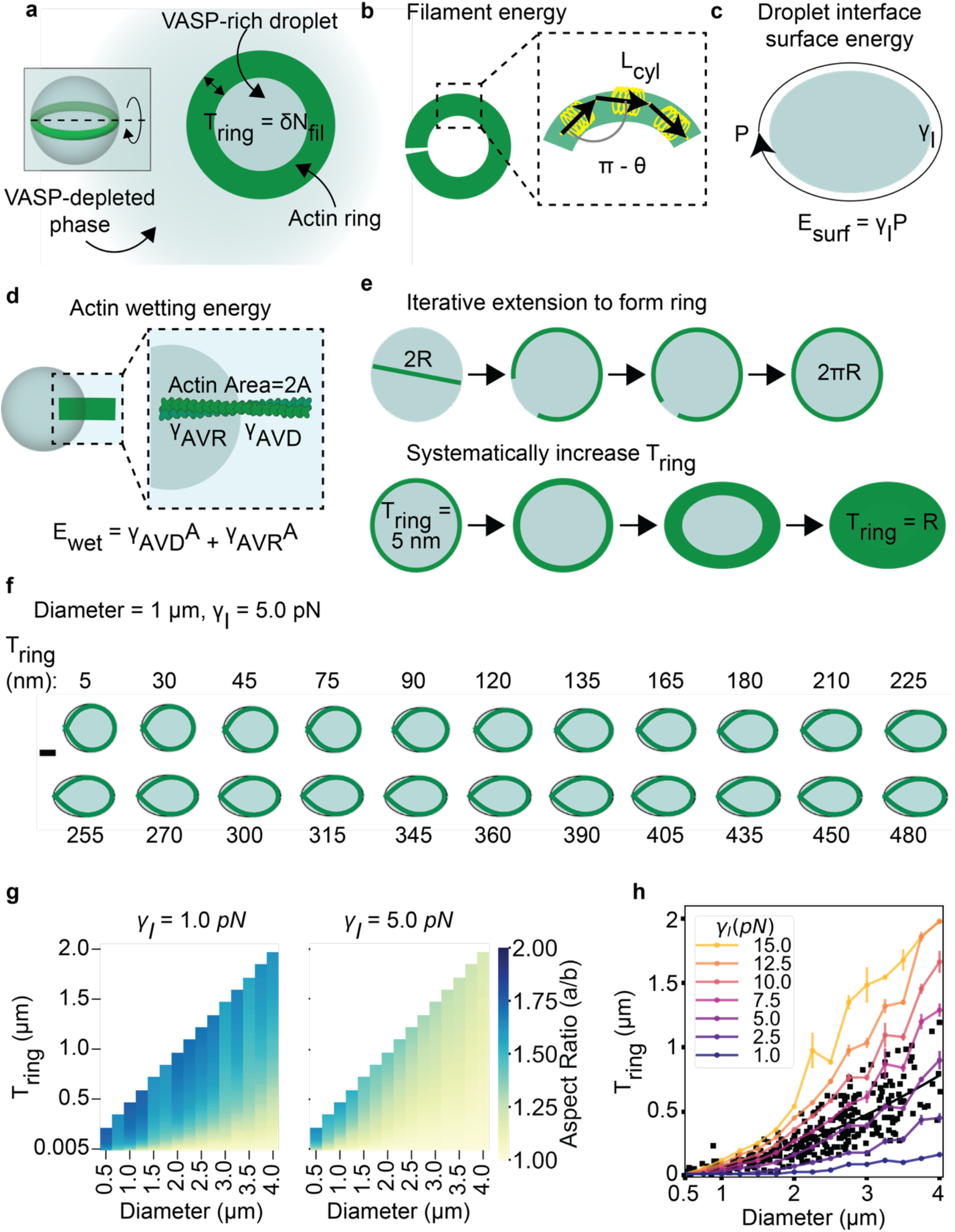
An elastic model predicts shape changes by balancing filament bending energy and droplet surface energy. (a) Cartoon shows a VASP-rich droplet phase in a VASP-depleted background. Droplet also contains an actin ring (green). We assume that the ring thickness scales linearly with the number of filaments. (b) Illustration of filament energy. Actin ring is represented as a linear chain discretized into segments of length L_cyl_ (stretching parameter, k_stretch_) that can bend around hinge points (parameter, k_bend_). (c) Illustration of VASP surface energy. The perimeter (P) of the VASP-rich droplet phase can change subject to surface energy parameter (*γ_I_*). (d) Illustration of actin wetting energy terms. Panel shows an actin ring of area 2A wetted by the VASP-dilute phase (AVD, interaction parameter, *γ_AVD_*) and VASP-rich phase (AVR, interaction parameter, *γ_AVR_*). Actin confinement to ring is obtained when *γ_AVR_* ≪ *γ_AVD_* as explained in Supplemental Meth©. (e) Cartoon shows the 2d plane of ring formation from panel a. Proposed 2D model accounts for shape changes due to actin deposition onto a preexisting ring. *Top:* Cartoon shows protocol to obtain an actin ring embedded in a 2d droplet from a droplet with single filament of length 2R through iterative increase of filament length in 100nm steps followed by energy minimization (energies explained in panels b-d). *Bottom:* Cartoon shows protocol to study shape changes at various ring thickness values. Starting with actin rings (as shown in panel c1), we generate minimum energy configurations of droplet-actin ring system through iterative increase of ring thickness (step size = 15nm) (f) Subpanels show the predicted shape of the droplet of initial radius 1 μm, and *γ_I_*=5.0pN when the actin ring (green line) thickness is increased from 5nm to 0.48μm. Scale bar = 0.25μm. Actin ring thickness visualized does not correspond to the Tring values used. (g) Heatmap shows the predicted aspect ratios of droplets of various initial diameters at various actin ring thicknesses and droplet diameter (*γ_I_*, =1.0 and 5.0 pN). (h) Plot shows the predicted mean and standard deviation (3 trials) ring thickness above which we observe the circle-ellipse shape transition (criterion: aspect ratio>1.1) at any given droplet diameter and droplet interfacial surface tension, *γ_I_*. Experimentally observed droplets from Fig 3e are overlaid as a scatter plot with black squares. Black solid line represents the trendline from a power law fit to the experimental data of the form y = a*x^b where a = 0.0721 and b = 1.7195 (R^2 = 0.6990).

### At higher actin to VASP ratios, droplets are elongated by actin polymerization

The computational model predicts that the aspect ratio of VASP droplets should increase with increasing stiffness of the actin rings that they encapsulate (Figure 4g,h). The validity of this prediction is directly demonstrated by the data in Figure 3f, showing that droplets with higher aspect ratios contain thicker, and therefore stiffer, actin rings. However, this analysis was confined to structures of relatively low aspect ratio (< 2.5) because increasing aspect ratio leads to narrowing of the droplet minor axis such that actin rings can no longer be resolved, Figure 2a (1:7.5). Nonetheless, the model in Figure 4 predicts that the aspect ratio of these structures should continue to increase with increasing actin content. To examine this idea, we plotted the distribution of droplet aspect ratios for each of the actin to VASP ratios examined in Figure 2a. As expected, we found that the median aspect ratio increased significantly with increasing actin to VASP ratio, Figure 5a.

**Figure 5:**
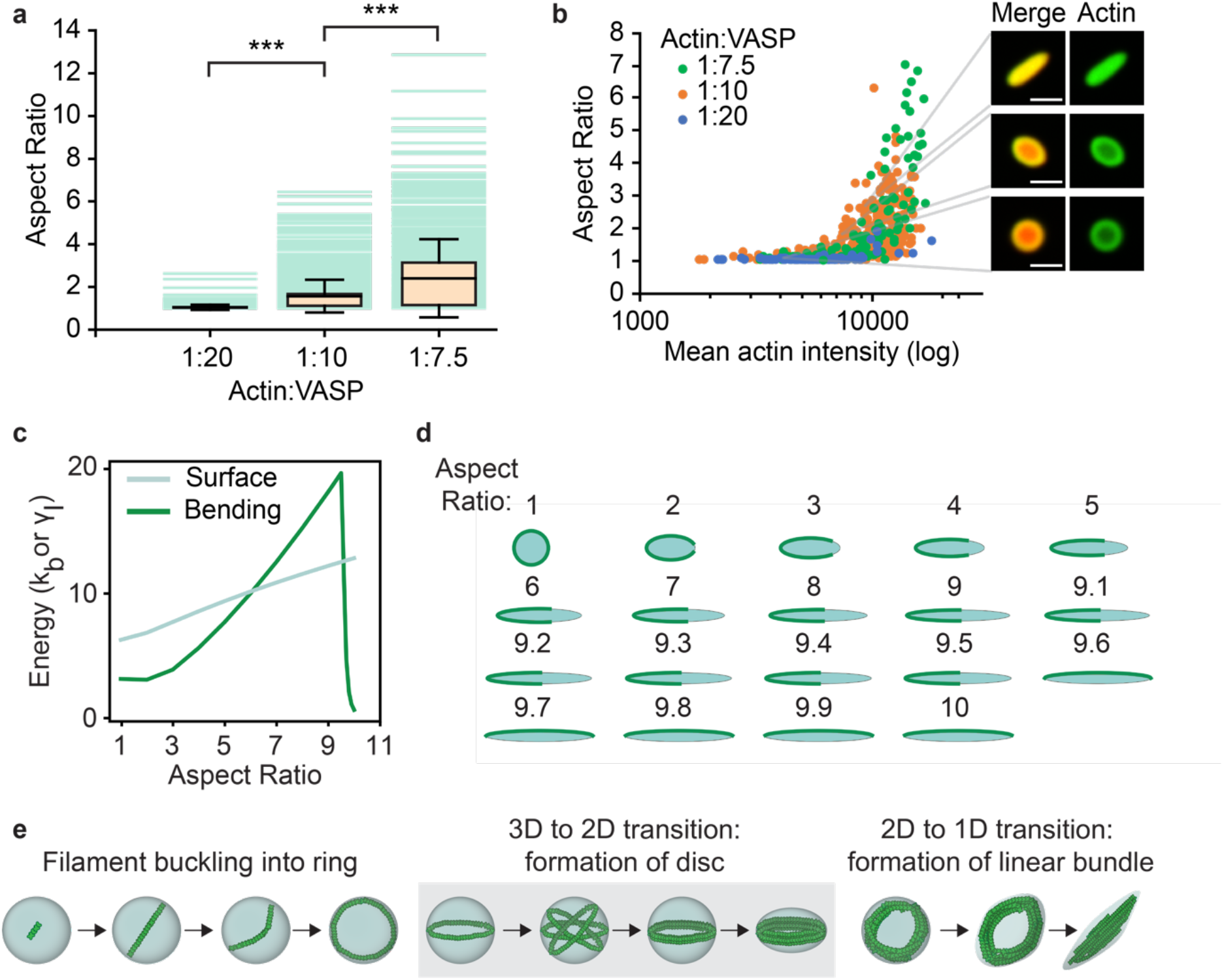
Increasing the actin concentration within the droplet drives droplet deformation into elongated structures. (a) Distribution of droplet aspect ratios observed across increasing actin to VASP ratios. Box represents interquartile range, with median as a bisecting line. Whiskers represent 1.5*SD of n = 496 droplets for 1:20, n = 1285 droplets for 1:10, and n = 589 droplets for 1:7.5 across 3 biologically independent experiments per condition. Each teal background line represents a single droplet. Brackets indicate data that was tested for significance using an unpaired, two-tailed t-test, with asterisks indicating p < 0.001. (b) Quantification of the dependence of droplet aspect ratio on actin intensity within the droplet (n = 626 droplets analyzed across at least 3 images per condition). Scale bar 2 μm. (c) Figure shows bundle bending energy (kb = 1pN.nm2) and droplet surface energy (γI = 1pN) at various droplet aspect ratios. Consider a droplet of radius R = D_droplet_/2 with a bundled-actin ring of contour length L = 2πR. The maximum permitted aspect ratio is obtained when 2a = L given ab = R2 resulting in (a/b)max = π2. As we increase aspect ratio of the droplet, the corresponding bending and surface energies are shown. (d) VASP droplet (blue) and bundled actin (green) are visualized at different aspect ratios (mentioned above). Bending energy increases as actin interacts with the high-curvature regions of the droplet. At larger aspect ratios (>9), minimum bending energy is obtained as droplet interacts entirely with the low-curvature regions of the droplet resulting in lower bending energy. Please refer to Supplementary Methods (Section 3.11) for detailed discussion of the mathematical model used. (e) Cartoon depicting hypothesized mechanism of droplet-driven actin bundling. Shaded region depicts droplets shown from 3D perspective, and unshaded regions depicts droplets shown from top-down perspective.

Examining the images in Figure 2a, it appears that droplets with higher actin content generally have higher aspect ratios. Variations in actin content likely arise from competition among droplets for actin, where droplets that initially sequester enough actin to drive polymerization attract more actin monomers, creating a spontaneous variation of actin intensities across the population of droplets. Notably, in the presence of latrunculin A, which prevents actin polymerization, the distribution of actin among VASP droplets was highly uniform, suggesting that actin polymerization gave rise to the variation in actin intensity, Figure 2. This variation presents an opportunity to examine the impact of actin concentration on droplet aspect ratio in situ. To study this effect, we plotted droplet aspect ratio as a function of actin intensity, Figure 5b and Supplemental Figure S8. Here, as expected, aspect ratio increased with actin intensity. Strikingly, data from different actin to VASP ratios overlay on a single trend, suggesting that actin concentration within the droplet is the dominant variable controlling aspect ratio, regardless of the bulk actin and VASP concentrations. Examining the shape of the curve in Figure 5b, the sharp rise in aspect ratio with increasing actin intensity suggests that there is a critical bundle rigidity, beyond which aspect ratio increases rapidly.

Once droplets take on elliptical shapes, what drives the transition into linear, rod-like structures? Using our continuum-scale model, this 2D to 1D transition can be understood as a further minimization of the bending energy of the filaments at the expense of droplet surface energy. A mechanical argument for this idea is provided in the Supplemental Material. Briefly, for a spherical droplet of circular cross-section, the radius of curvature is uniform everywhere and therefore, the bending energy is spatially homogeneous. However, as a droplet begins to elongate into an ellipsoid, it has an elliptical cross-section, which has regions of high curvature and regions of low curvature, Figure 5c,d. The disparity between the these curvatures increases sharply as the droplet aspect ratio increases, Figure 5d. Rather than accepting the increasingly high curvature at the ends of the ellipsoid, at a certain aspect ratio, it becomes energetically more favorable for the ring to straighten into a one-dimensional rod, Figure 5d.

In summary, by comparing experimental data to a continuum-scale computational model, we have shown that growing actin filaments deform initially spherical droplets of VASP through a series of symmetry breaking events, ultimately resulting in actin-filled droplets with rod-like shapes, Figure 5e.

### VASP droplets bundle actin filaments and exhibit liquid-like behavior

In cellular structures such as filopodia ^30^ and stress fibers ^31^, actin organizes into long bundles of very high aspect ratio. Therefore, we next investigated the impact of higher actin content on droplet aspect ratio. Specifically, we formed droplets from a solution of 10 μM VASP and added 2 μM of actin to achieve a 1:5 ratio of actin to VASP. Under these conditions, we observed rapid deformation of initially spherical droplets into thin, linear structures with aspect ratios that frequently exceeded 15:1, Figure 6a, b, Supplemental videos S5 and S6. Actin colocalized with VASP throughout the growth of these structures, suggesting that the droplets deformed and grew continuously with the growing actin filaments, Figure 6c. Do these structures consist of bundled, parallel actin filaments? If so, they should grow linearly and maintain a constant intensity in the actin channel over their length. The high aspect ratio structures in our experiments displayed these properties, with an average growth rate in the presence of ATP of 23 +/- 3 nm/s, about 8-9 monomers per second, Figure 6d. Figure 6a-c. This rate is several fold slower than the rates of VASP-catalyzed single filament elongation found in the literature ^1, 32^. The slower growth rate could arise from several factors including the viscosity of the droplet environment, which could slow the diffusion of new actin monomers to filament tips, and the possibility that not all filaments grow simultaneously, perhaps owing to competition for a limited supply of actin. In addition to a linear growth rate, bundles of parallel actin filaments should only grow through addition of new monomers to the tips of the bundle. In line with this prediction, when the tips of growing structures were bleached, the actin signal in the bleached region did not recover, as expected for stably polymerized filaments, Figure 6e,f (Supplemental Video S7). However, addition of fluorescent actin monomers, which must have originated from the surrounding solution, was observed at the filament tips, similar to the growth of parallel actin filaments inside filopodia^33 34^.

**Figure 6:**
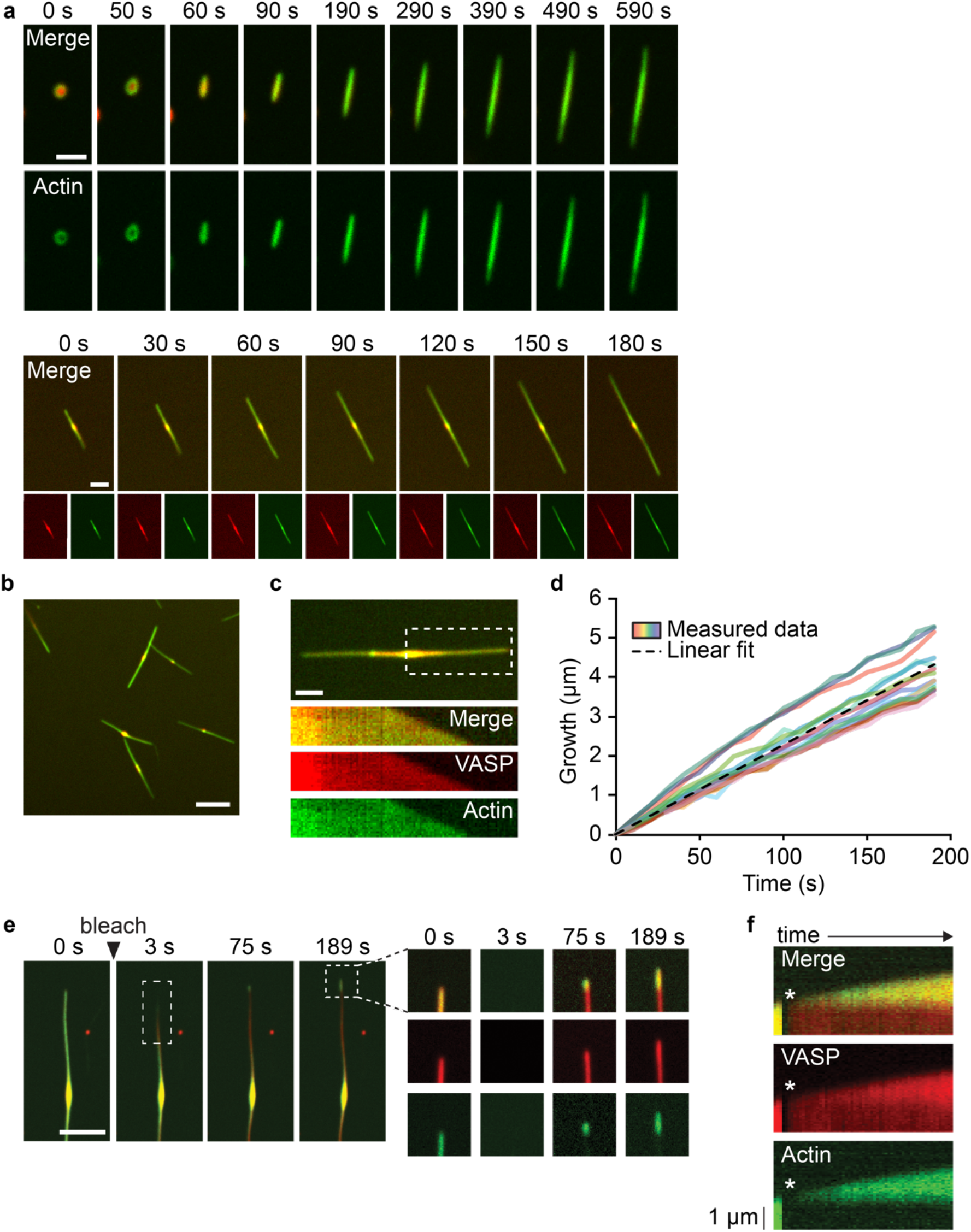
Linear droplets formed at high actin to VASP ratios consist of parallel bundled actin filaments. (a) Addition of 2 μM actin added to 10 μM VASP droplets results in linear droplets that elongate over time. Scale bar 5 μm. (b) Representative image of linear droplets generated after adding actin to VASP droplets. Scale bar 5 μm. (c) Representative image (top) and associated kymograph (bottom) of a linear droplet elongating over time. Scale bar 2 μm. (d) Quantification of the elongation rates of linear droplets (n= 15 droplets over 3 biologically independent experiments). Trendline represents the linear fit of the average across all droplets. Linear fit y = 0.0227x R^2^ = 0.9965. (e) Photobleaching of the ends of the elongating droplets results in recovery at the tips of the bundle. Dotted white box indicates bleach region. Scale bar 2 μm. (f) Kymograph of the recovery profile of the droplet bleached in E. Asterisk denotes the bleach frame.

In contrast to actin, the VASP signal recovered within minutes, suggesting that the VASP phase remained liquid-like. Additionally, when growing actin bundles encountered one another, they “zippered” together within seconds, presumably to minimize the surface energy of the liquid-like VASP droplets in which they were confined, Figure 7a, Supplemental Video S8. Is a fluid-like VASP matrix required for actin bundling? To further elucidate the role of droplet fluidity in actin bundling, we tested whether more solid droplets were capable of bundling actin. Specifically, when we increased the concentration of PEG from 3% to 10%, the diameter of VASP droplets remained approximately the same (Supplemental figure S9), but the droplets became solid-like, as demonstrated by very slow rates of recovery in FRAP experiments (Figure 7b), and a marked decrease in fusion events (Supplemental figure S9). In the presence of actin, the fraction of droplets that deformed into rods decreased with increasing PEG concentration (Figure 7c, Supplemental figure S9). Specifically, with 10% PEG, a majority of the droplets remained spherical, whereas the majority of droplets at 3% PEG transformed into rods (Figure 7d, Supplemental figure S9). Collectively, these observations suggest that the liquid-like nature of VASP droplets is essential to bundling of actin filaments.

**Figure 7:**
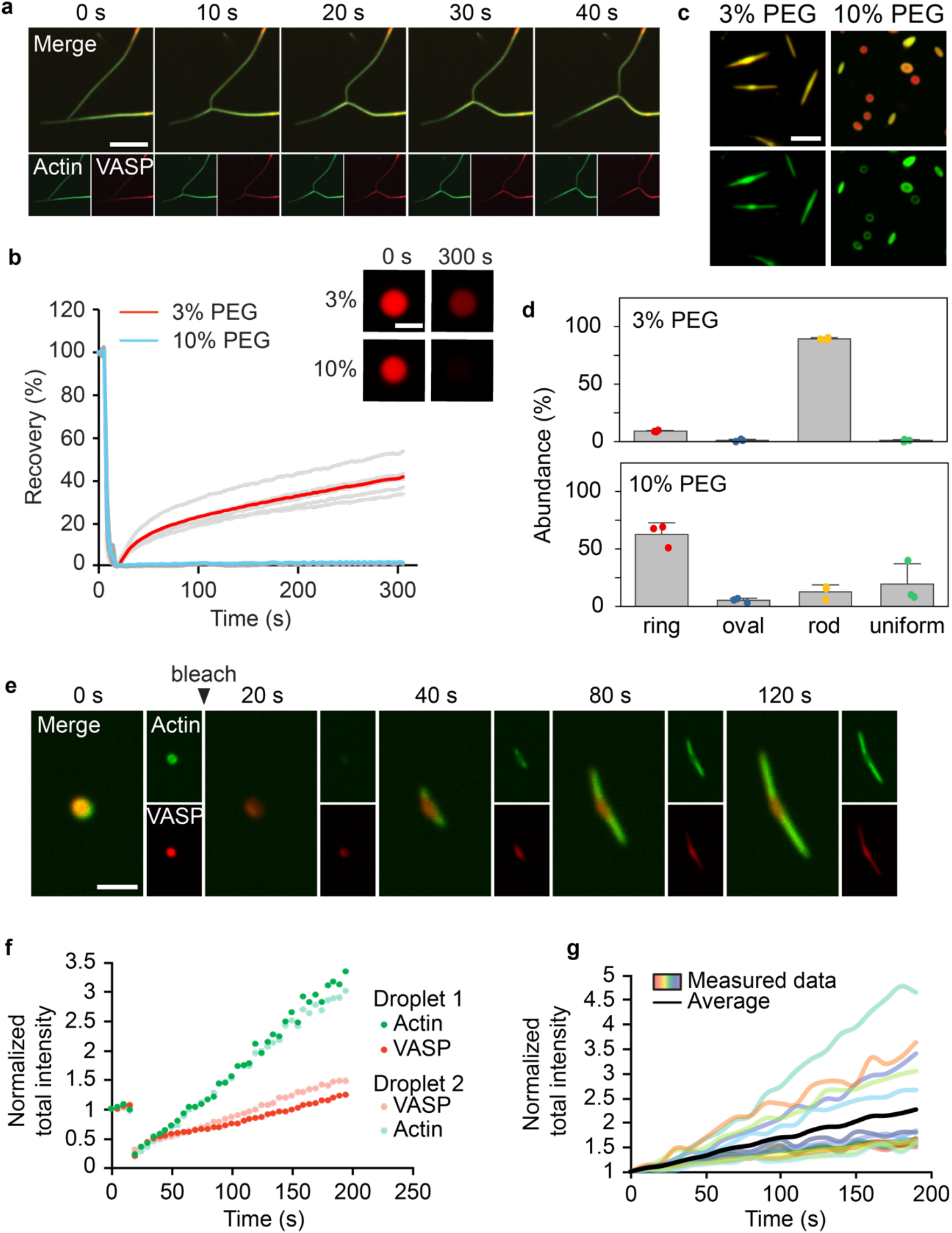
Liquid-like behavior is required for robust actin bundling. (a) Droplet bundles that come in contact zipper together to form a thicker bundle. Scale bar 5 μm. (b) 10 μM VASP droplets formed at increasing PEG concentrations are less mobile, as measured by FRAP. Error bars represent SD across n = 5 droplets. *Inset:* Images depicting droplets formed at 10% PEG recover to a lesser extent than droplets formed at 3% PEG. Scale bar 2 μm. Note, recovery rates are slower than those in Figure 1f because a larger area was bleached. (c) 10 μM VASP droplets (red) formed in droplet buffer containing 10% PEG deform from 2 μM actin to a lesser extent than at 3% PEG. Scale bar 5 μm. (d) 10 μM VASP droplets formed at 10% PEG result in less rod formation after polymerization of 2 μM actin. Bars represent mean and whiskers represent standard deviation across n = 3 biologically independent experiments. At least 500 droplets were analyzed per condition. (e) Photobleaching of a round droplet that subsequently undergoes transformation into a rod. Scale bar 2 μm. (f) Quantification of the total actin and VASP intensity in the structure over time, as seen in E. Total actin intensity is the result of summing the pixel intensities across the droplet and normalizing to the pre-bleach total intensity. (g) Quantification of the total actin intensity in elongating VASP bundles from 6d over time. Total actin intensity is the result of summing the pixel intensities across the droplet and normalizing to the total intensity of the droplet at the first time point.

Liquid-like droplets should permit additional molecules of actin and VASP to enter growing structures. To probe this effect, we performed FRAP on spherical droplets, just as they were beginning to transform into bundles, Figure 7e and Supplemental Video S9. As these droplets deformed, the total actin intensity increased dramatically during growth of the bundle, greatly exceeding the initial actin intensity and dimensions of the droplet before it was bleached, Figure 7f. Meanwhile, the intensity in the VASP channel also exceeded its initial value, albeit more modestly. Similar results were observed for elongating, unbleached bundles, Figure 7g. These results demonstrate the ability of the condensed VASP phase to template filament alignment. Once growing actin filaments become aligned within a VASP droplet, the parallel orientation of filaments propagates during the growth of the bundle as new material from outside the initial droplet is incorporated. In this way, even a small droplet of VASP can setup a parallel orientation of actin filaments that will persist over much longer length scales.

## DISCUSSION

Here we show that the actin polymerase, VASP, assembles into liquid-like droplets that polymerize and bundle actin. Polymerization of actin inside these droplets drives a series of morphological transformations from spherical droplets with shell and ring-like actin distributions to deformation of the surrounding droplets into discs, ellipsoids, rods, and finally linear bundles of parallel filaments. These transformations can be understood by minimizing the energy of the filament-droplet composite, as illustrated by a continuum-scale computational model.

Previous work has shown that non-specific interactions between actin and the surfaces of protein droplets can concentrate actin, driving its polymerization outside of droplets ^35^, and that incorporation of short, pre-polymerized filaments can promote filament alignment, leading to nematic crystals with tactoid-like droplet morphologies ^36^. In contrast, our work illustrates the potential of droplets to couple actin polymerization to alignment of filaments, ultimately leading to droplet-encapsulated filament bundles that are self sustaining. Earlier studies illustrated that droplets of the tau protein can nucleate microtubule polymerization, ultimately leading to filament alignment ^37^. However, microtubules have persistence lengths on the millimeter scale ^38^, 100-fold larger than the persistence length of actin. Therefore, microtubules are far too rigid to organize within droplets by the mechanisms that we have illustrated for actin filaments.

Our primary finding is that liquid-like protein droplets can mediate the bundling of semi-flexible actin filaments through a mechanism in which the rigidity of a growing filament bundle competes against the finite surface energy of the droplet. Further, our data illustrate how a droplet can locally concentrate actin, and promote polymerization through continuous flux of new monomers. Together, these features lead to spontaneous organization of the growing actin filaments into a ring-like bundle. Catalyzed by the high concentration of actin and VASP in the droplet, additional actin assembly results in a thickening of the actin ring until its bending rigidity overcomes the surface tension of the droplet. Once sufficiently rigid, the ring deforms the droplet, transforming it into a linear structure that grows with a constant polymerization rate. Zippering together of bundles upon contact, an effect driven by the liquid-like nature of VASP, further increases the extent of filament bundling.

VASP associates with aligned actin filaments in multiple cellular structures including filopodia and focal adhesions ^39, 40^. Interestingly, multiple proteins involved in assembly of focal adhesions have recently been found to undergo liquid-liquid phase separation ^41–43^. However, the role of this network in aligning actin filaments is presently unknown. Our work builds on this understanding by demonstrating a new physical mechanism by which VASP, and potentially other actin interacting proteins, can harness phase separation to drive spontaneous, self-sustaining growth of actin bundles.

## MATERIALS & METHODS

### Reagents

Tris base, NaCl, EDTA, EGTA, TCEP, poly-L-lysine, Atto 594 maleimide, catalase, glucose oxidase were purchased from Sigma-Aldrich. Alexa Fluor 647 C2 maleimide was purchased from Thermo Fisher Scientific. Phalloidin-iFluor488 was purchased from Abcam. Amine-reactive PEG (mPEG–succinimidyl valerate MW 5000) was purchased from Laysan Bio. Rabbit muscle actin was purchased from Cytoskeleton.

### Cloning and protein purification

A pET vector encoding the “cysteine light” variant of human VASP (pET-6xHis-TEV-KCK-VASP(CCC-SSA)) was a generous gift from Scott Hansen. All VASP mutants were generated using this plasmid as a template. pET-6xHis-TEV-KCK-VASPΔtet encoding monomeric VASP was generated by using site directed mutagenesis to introduce a stop codon after residue 339 to generate a truncated version of VASP lacking residues 340-380 corresponding to the tetramerization domain. The forward primer: “GCTCCAGTTAGTACTCGGACCTACAGAGGG” and reverse primer “GTCCGAGTACTAACTGGAGCTGGGCGTG. pET-6xHis-TEV-KCK-VASPΔEVH1 encoding VASPΔEVH1 was generated by PCR mediated deletion of the EVH1 domain (residues 1-113). The 750 base pairs corresponding to the EVH1 domain were deleted using the forward primer: “GACTAAGCGGCCGCGAAGGAGGTGGGCCCC” and reverse primer: “GACTAAGCGGCCGCCTTTACATTTGGATCCCTGGAAGTACAG” that both encoded NotI cut sites. After PCR amplification and purification, the amplicon was digested with NotI and recircularized to generate the resulting product. pGEX-6P1-KCK-IDR encoding the intrinsically disordered, proline-rich region of VASP was generated by restriction cloning. Amplification of the IDR (residues 115-222) was achieved using the forward primer: “CAACGAGGATCCAAATGTAAAGGAGGTGGGCCCCCTCCAC” and reverse primer: “TAATATCTCGAGTAATTACCCAGCTCCCCCACCACCAGGG”. The amplicon was then inserted into the pGEX-6P1 vector at the BamHI and XhoI restriction sites.

The pET-His-KCK-VASP(CCC-SSA) plasmid was transformed into *E. coli* BL21(DE3) competent cells (NEB Cat#C2527). Cells were grown at 30°C to an OD of 0.8. Protein expression was performed as described previously with some alteration ^1^. Expression of VASP was induced with 0.5mM IPTG, and cells were shaken at 200 rpm at 12°C for 24 hours. The rest of the protocol was carried out at 4°C. Cells were pelleted from 2L cultures by centrifugation at 4785 x g (5000 rpm in Beckman JLA-8.100) for 20 min. Cells were resuspended in 100ml lysis buffer (50mM sodium phosphate pH 8.0, 300mM NaCl, 5% Glycerol, 0.5mM TCEP, 10mM imidazole, 1mM PMSF) plus EDTA free protease inhibitor tablets (1 tablet/50ml, Roche cat#05056489001), 0.5% Triton-X100, followed by homogenization with a dounce homogenizer and sonication (4×2000J). The lysate was clarified by ultracentrifugation at 125,171 x g (40K rpm in Beckman Ti45) for 30 min. The clarified lysate was then applied to a 10ml bed volume Ni-NTA agarose (Qiagen Cat#30230) column, washed with 10xCV of lysis buffer plus EDTA free protease inhibitor tablets (1 tablet/50ml), 20mM imidazole, 0.2% Triton X-100, followed by washing with 5xCV of lysis buffer plus 20mM imidazole. The protein was eluted with elution buffer (50mM Tris, pH 8.0, 300mM NaCl, 5% Glycerol, 250mM imidazole, 0.5mM TECP, EDTA free protease inhibitor tablets (1 tablet/50ml)). The His-tag was cleaved by dialyzing the protein overnight with 1mg (10,000 units) of TEV protease in 1L of cleavage buffer (50mM Tris pH 8.0, 200mM NaCl, 5% Glycerol, 0.5mM EDTA, 1mM DTT). The His-tag, uncut protein, and TEV protease were removed by a second Ni-NTA column. The protein was further purified by size exclusion chromatography with Superose 6 resin. The resulting purified KCK-VASP was eluted in storage buffer (25mM HEPES pH 7.5, 200mM NaCl, 5% Glycerol, 1mM EDTA, 5mM DTT). Single-use aliquots were flash-frozen using liquid nitrogen and stored at −80°C until the day of an experiment. The his-tagged VASP mutants were purified using the same protocol as above as indicated or with the following modifications: His-KCK-VASPΔTet: no modifications. His-KCK-VASP-CRY2: the pH of all the buffers was changed to 8.5.

The pGEX-GST-KCK-VASP-PRR plasmid was transformed into *E. coli* BL21 competent cells (NEB Cat#C2530). Cells were grown at 30°C to an OD of 0.8. Protein expression was induced with 0.9mM IPTG, and cells were shaken at 200 rpm at 30°C for 6-8 hours. The rest of the protocol was carried out at 4°C. The cells were pelleted from 2L cultures by centrifugation at 4785 x g (5000 rpm in Beckman JLA-8.100) for 20 min. Cells were resuspended in 100ml lysis buffer (20mM Tris pH 8.0, 350mM NaCl, 10mM KCl, 5% Glycerol, 5mM EDTA, 5mM DTT, 1mM PMSF) plus EDTA free protease inhibitor tablets (1 tablet/50ml, Roche cat#05056489001), 1.0% Triton-X100, followed by homogenization with a dounce homogenizer and sonication (4×2000J). The lysate was clarified by ultracentrifugation at 125,171 x g (40K rpm in Beckman Ti45) for 30 min. The clarified lysate was then applied to a 10ml bed volume Glutathione Sepharose 4B (Cytiva Cat#17075605) column, washed with 10xCV of lysis buffer plus 0.2% Triton X-100, EDTA free protease inhibitor tablets (1 tablet/50ml), followed by washing with 5xCV of lysis buffer. The protein was eluted with lysis buffer plus 15mM reduced glutathione, EDTA free protease inhibitor tablets (1 tablet/50ml). The GST-tag was cleaved by exchanging the protein into 20mM Tris pH 8.0, 350mM NaCl, 10mM KCl, 5% Glycerol, 5mM EDTA, 1mM DTT using a Zeba desalting column (Thermo Scientific cat#89891), adding PreScission protease (Thermo Scientific: cat 88946, 2unit/0.2mg fusion protein), and rocking gently overnight at 4°C. The GST-tag, uncut protein, and PreScission protease were removed by a second Glutathione Sepharose 4B column. The protein was further purified by size exclusion chromatography with Superose 6 resin. The resulting purified KCK-VASP was eluted in storage buffer (20mM Tris, pH 8.0, 200mM NaCl, 5% Glycerol, 1mM EDTA, and 5mM DTT). Single-use aliquots were flash-frozen using liquid nitrogen and stored at −80°C until the day of an experiment.

GST-KCK-VASPΔEVH1 was purified using the same protocol as above, but with the following buffer modifications: Lysis Buffer=20mM Tris pH 8.5, 350mM NaCl, 5% Glycerol, 5mM EDTA, 5mM DTT, 1mM PMSF; Storage Buffer=25mM HEPES, pH 7.5, 200mM NaCl, 5% Glycerol, 1mM EDTA, and 5mM DTT.

### Protein labeling

We utilized a previously published mutant of human VASP, in which the three endogenous cysteines were replaced by two serines and an alanine, respectively. A single, exogenous cysteine was introduced at the N-terminus of the protein allowing for selective labeling using maleimide chemistry ^1^. This mutant has been found to drive actin polymerization at rates indistinguishable from the wild-type protein ^1^.

VASP and associated VASP mutants were selectively labeled at the single N-terminal cysteine residue using maleimide conjugated dyes. Protein was incubated with a 3-fold molar excess of dye for 2 hours at room temperature to achieve a 10-20% labeling ratio. The protein was separated from the free dye by applying the labeling reaction to a Princeton CentriSpin size exclusion column (Princeton Separations). Labeled VASP VASPΔtet, and VASPΔEVH1 were stored in 50 mM Tris pH 7.8, 300 mM NaCl, 0.5 mM EDTA 0.5 mM EGTA, 5 mM TCEP, and VASP-IDR was stored in 25 mM HEPES pH 7.1 300 mM NaCl 0.5 mM EDTA 0.5 mM EGTA 5 mM TCEP. Monomeric actin was labeled at Cys-374 using maleimide conjugated dyes. Dye was incubated with G-actin at a 3-5 fold molar excess for 2 hours at room temperature to achieve a labeling ratio of 20-30%. Unconjugated dye was separated from labeled actin by applying the labeling reaction to a Princeton CentriSpin-20 size exclusion column hydrated with A buffer (5 mM Tris-HCl pH 8, 0.2 mM CaCl2, 0.2 mM ATP, 0.5 mM DTT). The eluted labeled protein was then centrifuged at 14k rpm for 15 min at 4°C to remove aggregates, and flash-frozen in singleuse aliquots.

### VASP droplet formation and actin polymerization

Unless otherwise noted, VASP droplets were formed by mixing VASP with 3% (w/v) PEG-8000 in 50 mM Tris pH 7.4 150 mM NaCl 0.5 mM EDTA 0.5 mM EGTA 5 mM TCEP. PEG was added last to induce droplet formation. All VASP concentrations refer to the concentration of VASP monomer.

For actin polymerization assays within VASP droplets, VASP droplets were formed and monomeric actin was mixed into the droplet solution. For end point actin assays (Fig 2a-d, Fig 5), Atto-488 labeled G-actin was allowed to polymerize in the droplets for 15 minutes, then the droplets were imaged. An actin to VASP ratio of 1:20 corresponds to 20 μM VASP and 1 μM actin, 1:10 corresponds to 20 μM VASP and 2 μM actin, and 1:7.5 corresponds to 15 μM VASP and 2 μM actin. For phalloidin-actin assays (Fig 3a, c-f), unlabeled actin monomers were added to VASP droplets and allowed to polymerize for 15 minutes. Phalloidin-iFluor488 (Abcam) was added to stain filamentous actin for 15 minutes, then the droplets were imaged. For quantification of phalloidin-stained ring thicknesses, 20 μM VASP droplets were formed and 1, 2, or 2.67 μM actin was added to yield actin to VASP ratios of 1:20, 1:10, and 1:7.5. For time lapses of actin polymerization within VASP droplets (Fig 2 e-f, Figs 6 and 7), VASP droplets were prepared in droplet buffer supplemented with oxygen scavengers (130 μg/ml glucose oxidase, 24 μg/ml catalase, and 40 mM glucose), actin-488 was added, and imaging was initiated within 3 minutes. For quantification of growing actin bundles (Figures 6 and 7) droplet buffer without EDTA or EGTA, and supplemented with scavengers and 1 mM ATP was used.

For FRAP (Fig 1f) 15 uM VASP droplets labeled with 3% Atto-488 were formed as above, in droplet buffer supplemented with oxygen scavengers. A 1.8 μm region within the droplet was bleached. For FRAP of 10 μM VASP droplets formed with increasing PEG, droplets were supplemented with oxygen scavengers and whole droplets of 2 μm diameter were bleached.

### Microscopy

Samples were prepared in wells formed by 1.5 mm thick silicone gaskets (Grace Biolabs) on Hellmanex II (Hellma) cleaned, no.1.5 glass coverslips (VWR) passivated with PLL-PEG. A second coverslip was placed on top to seal the imaging chamber to prevent evaporation during imaging. Fluorescence microscopy was performed using the Olympus SpinSR10 spinning disc confocal microscope fitted with a Hamamatsu Orca Flash 4.0V3 SCMOS Digital Camera. FRAP was performed using the Olympus FRAP unit 405 nm laser. In some cases, super resolution with deconvolution was performed to better visualize actin filaments within VASP droplets.

PLL-PEG was prepared as described previously with minor alteration ^44^. Briefly, amine-reactive mPEG-SVA was conjugated to Poly-L-Lysine (15-30 kD) at a molar ratio of 1:5 PEG to poly-L-lysine. The conjugation reaction was performed in 50 mM sodium tetraborate pH 8.5 solution and allowed to react overnight at room temperature with continued stirring. The product was buffer exchanged into PBS pH 7.4 using Zeba spin desalting columns (7K MWCO, ThermoFisher) and stored at 4°C.

### Image analysis

ImageJ was used to quantify the distribution of droplet characteristics. Droplets were selected by thresholding in the VASP channel, and shape descriptors (e.g. diameter, aspect ratio) and protein intensities were measured using the analyze particles function. Quantification of actin shapes in droplets was performed by scoring 3 images from each replicate, (n= 3 total replicates) for the presence or absence of uniform actin, rings, ovals, and rods. Uniform droplets were defined as droplets with AR <1.1 that contain uniform actin staining. Rings were defined as droplets with AR < 1.1 that exhibited a decrease in fluorescence intensity in the center of the droplet. Ovals were defined as droplets with AR > 1.1 that exhibited a decrease in fluorescence intensity in the center of the droplet. Rods were defined as droplets with AR > 1.1 that contain uniform actin staining.

Quantification of the ring thickness within the droplets was performed by measuring the phalloidin-actin intensity across a radial line bisecting the ring, and taking the full width at half maximum intensity (with the diffraction limit subtracted). In order to compare droplet diameters among aspherical droplets with AR > 1, we calculated an effective spherical diameter for the droplets based on their major and minor axis lengths: 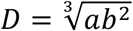 where a refers to the major axis and b refers to the minor axis lengths. Rings, discs, and shells were differentiated by making 3D projections of each individual droplet. Shells were defined as spherical droplets that have uniform actin intensity in the 3D projection. Rings were defined as spherical droplets that contained a planar arrangement of actin in the 3D projection. Discs were defined as non-spherical, planar droplets that colocalized both actin and VASP. Only droplets with peripheral accumulation of actin in the imaging plane were counted.

To calculate the elongation rate of bundles in ImageJ, single droplets were selected by thresholding, and the long axis length was measured over time using the analyze particles function. The change in length per frame (termed “growth”) was then plotted against time, and the slope of the line was used to determine the elongation rate. Only droplets that elongated into straight linear bundles, and persisted for at least 10 frames were counted, ie: bundles that curved, or elongated out of the focal plane were discarded. Bundles that were a product of multiple droplets merging or zippering were also discarded. For bundles that elongated bidirectionally, the droplet was segmented in the middle origin and the two elongating halves were measured independently. To calculate the total intensity of actin within the growing bundles, single droplets were selected by thresholding in the VASP channel, and the corresponding total intensity in the actin channel was measured.

FRAP data was analyzed using the FRAP Profiler plugin for ImageJ. Fluorescence recovery of a 1.8 μm diameter region was measured over time, and intensities were normalized to the maximum pre-bleach and minimum post-bleach intensity. All intensity data was corrected for photobleaching. Recovery was measured only for droplets of similar diameters.

## Acknowledgements

This research was supported by grants from the National Institutes of Health to JCS (R35GM139531) and PR (R01GM132106), by the National Science Foundation through a Modulus Grant BIO-1934509 to PR and JCS, and by the Welch Foundation through Grant F-2047 to JCS. The authors would like to thank Prof. Sapun Parekh and Daniel Dickinson for their feedback. AC would like to thank Dr. Ashwin Ravichandran for thoughtful discussions on the model. The authors declare no competing interests.

## Data availability

All data that support this work are available from the corresponding author upon request.

## Code availability

ImageJ (version 2.1.0) was used with the FRAP profiler plugin, which is available online (https://worms.zoology.wisc.edu/ImageJ/FRAP_Profiler_v2.java). No original code was generated in this work.

