## Supplemental Material for "Liquid-like VASP condensates drive actin polymerization and dynamic bundling"

### Supporting Information: Liquid-like assembly of VASP drives actin polymerization and bundling

### 1 Supplementary Figures

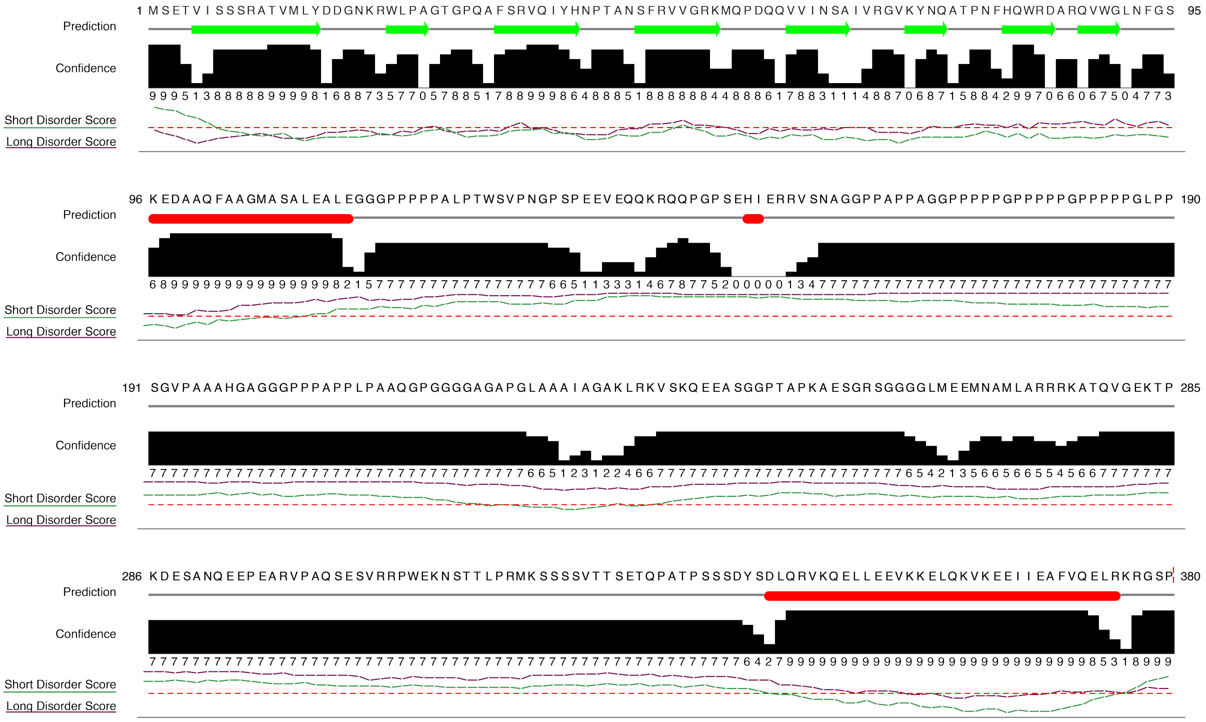

Supplemental Figure S 1: **Structural and disorder prediction of VASP by Jpred and IUPred.** Results from Jpred secondary structure prediction [1]. Prediction reveals the secondary structure prediction, with green arrows representing predicted sheets, and red ovals representing predicted helices. The confidence estimate of the prediction is noted on a scale from 0-9. Results from IUPred long disorder score predicts disordered domains of at least 30 consecutive residues, while short disorder score predicts shorter disordered regions [2, 3]. Results are depicted as a score from 0 to 1, with scores above 0.5 (red dotted line) indicate a higher probability of being disordered. Alignment of sequence predictions was achieved using Jalview [4].

.

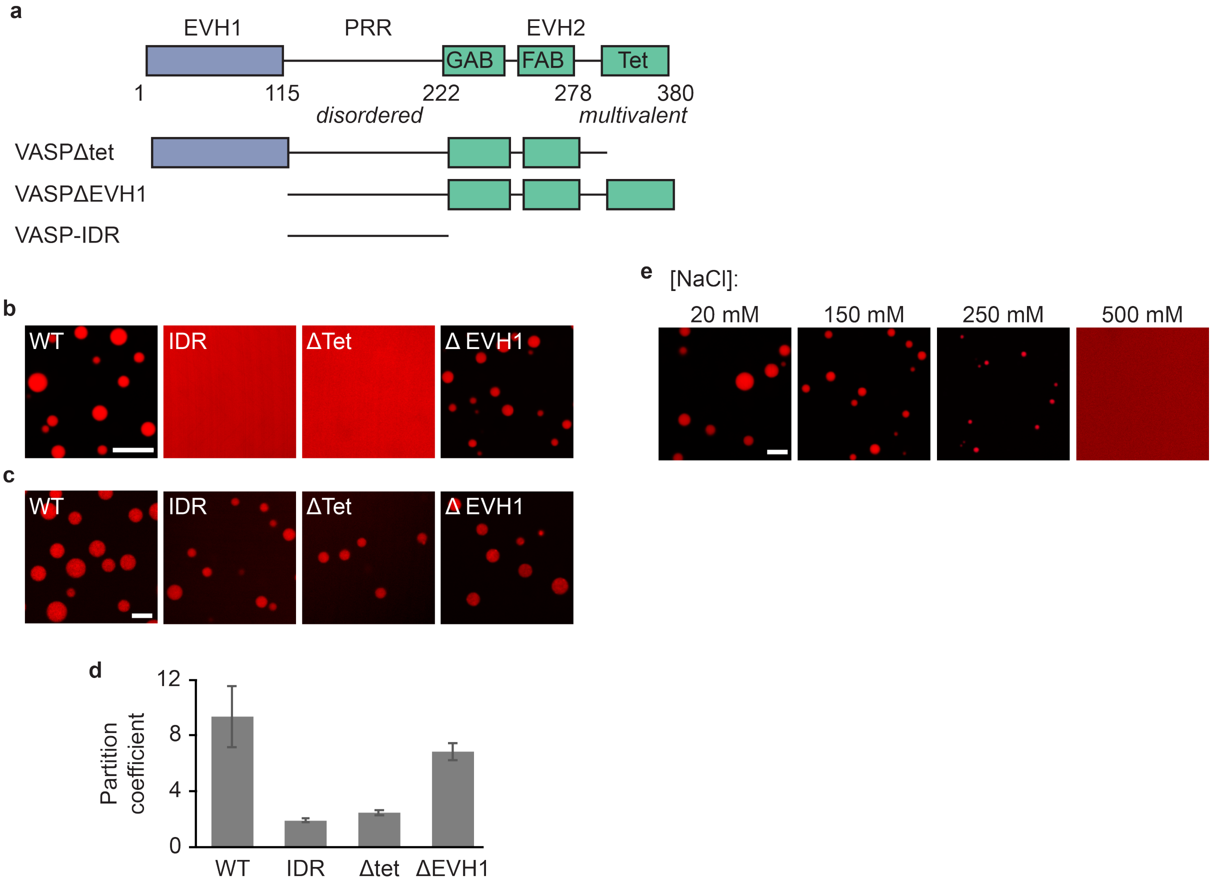


Supplemental Figure S 2: **VASP droplet formation relies on weak electrostatic interactions between tetramers.** (a) Schematic of VASP domain organization and designed mutants. (b) Droplet formation tests of various VASP mutants. Since intrinsically disordered regions are in many cases sufficient to drive LLPS[5], we tested whether the intrinsically-disordered, proline-rich region of VASP (VASP-IDR) was capable of forming droplets on its own. We found that VASP-IDR did not form droplets, even at a protein concentration of 100 $\mu M$ and a PEG concentration of $10\%$ (w/v). Since multivalency is critical for generating the higher order networks required for LLPS, we next tested whether monomeric VASP, which lacks the coiled-coil domain responsible for tetramerization, VASPΔtet, was capable of forming droplets. VASPΔtet was unable to form droplets, even at high protein or PEG concentrations (up to 56 $\mu M$ VASP and up to $5\%$ PEG). We next sought to determine how tetramers associate to generate an extended network. None of the domains within VASP are known to bind to one another, suggesting that the interactions between tetramers are weak. One possibility is that the EVH1 domain, which is known to interact with specific proline repeat sequences[6], could interact with VASP's proline-rich IDR. However, the specific proline-motifs to which EVH1 binds are not present in the IDR, suggesting that these interactions would be very weak. Nonetheless, we constructed a mutant of VASP with the EVH1 domain deleted, VASPΔEVH1. This mutant displayed significantly impaired phase separation, as 30 $\mu M$ VASPΔEVH1 was unable to form droplets at $3\%$ PEG (data not shown), but formed small droplets at $5\%$ PEG (data not shown), and larger ones when the protein concentration increased to 100 $\mu M$, while maintaining $5\%$ PEG. These data suggest that the EVH1 domain plays a role in LLPS of VASP, perhaps by interacting with VASP's IDR, though the exact interaction remains unclear. Scale bar 10 $\mu m$. (c) Partitioning of 1 $\mu M$ VASP and VASP mutants to 20 $\mu M$ unlabeled VASP droplets. Scale bar 5 $\mu m$. (d) Quantification of partitioning of 1 $\mu M$ of each of the mutants into droplets formed from 20 $\mu M$ of full-length VASP. Partitioning is defined as the ratio of protein intensity inside the droplet to outside the droplet. Bars depict averages, and error bars represent standard error across n=3 independent experiments with at least 3 images quantified per experiment. (e) Increasing salt concentration disrupts 15 $\mu M$ VASP droplet formation, indicating that electrostatic interactions are important for VASP condensation. This finding is consistent with the high density of charged residues in the EVH1 and EVH2 domains of VASP, which may result in weak electrostatic interactions that help create a long-range network[7–9]. Scale bar 5 $\mu m$.

.

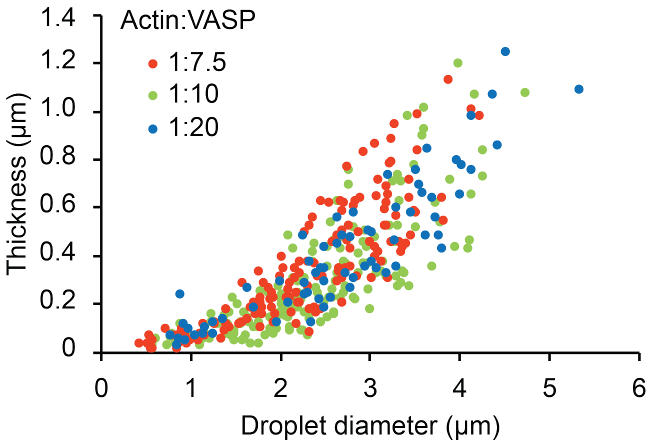


Supplemental Figure S 3: **Ring thickness is not determined by solution actin to VASP ratio.** Quantification of ring thickness as a function of droplet diameter across the three actin to VASP ratios tested (n = 386 droplets). All droplets are for aspect ratios less than 1.1.

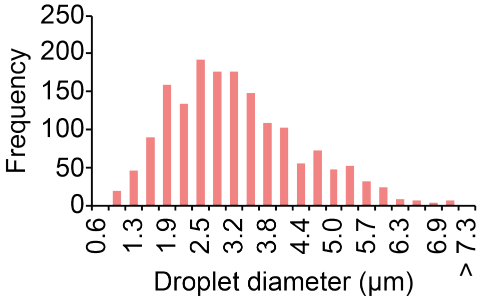


Supplemental Figure S 4: **Histogram of the distributions of droplet sizes for 20 $\mu M$ VASP droplets.** Median droplet size is 2.86 $\mu m$. Maximum droplet size is 7.22 $\mu m$. n= 1656 droplets counted over at least 3 images from 3 independent replicates.

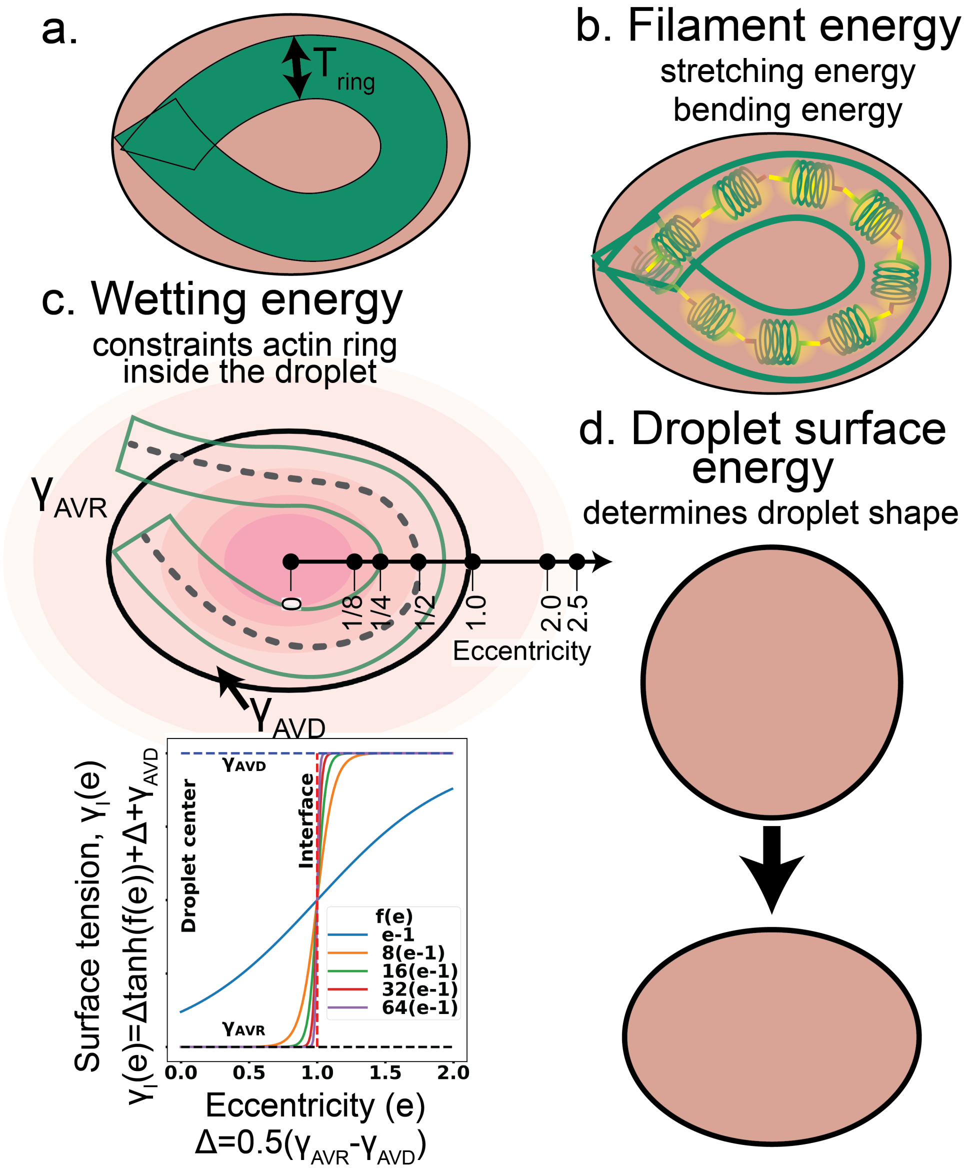


Supplemental Figure S 5: **Energetic considerations to account for the shape change of a VASP droplet containing a bundled-actin ring.** a. A cartoon of a bundled-actin ring of thickness $T_{ring}$ is shown in a VASP-rich droplet. The interface between the VASP-rich and -depleted phases is shown in black. The total energy of the system is obtained by adding the following terms b. **Filament energy** – we represent the actin bundle as a linear polymer discretized into segments of size 100 nm. We consider the bundled-actin ring to be made of multiple actin filaments spaced 15 nm apart. The persistence length of the ring is assumed to scale linearly with the number of filaments. The total energy in the bundled-actin ring is given by the sum of stretching and bending energy. We assume that the bundled-actin ring thickness does not change during the structural transitions. c.**Wetting energy** – Interaction energy parameters $(\gamma_{AVR} and \gamma_{AVD})$ are chosen to ensure actin prefers to interact with the VASP-rich droplet. Interaction parameter at the interface is defined by a hyperbolic tangent function to ensure continuity and differentiability. d. **Surface energy** – While the total surface area of the droplet is held constant during the simulation, the energetic cost of changing droplet perimeter is given by $\gamma_I$.

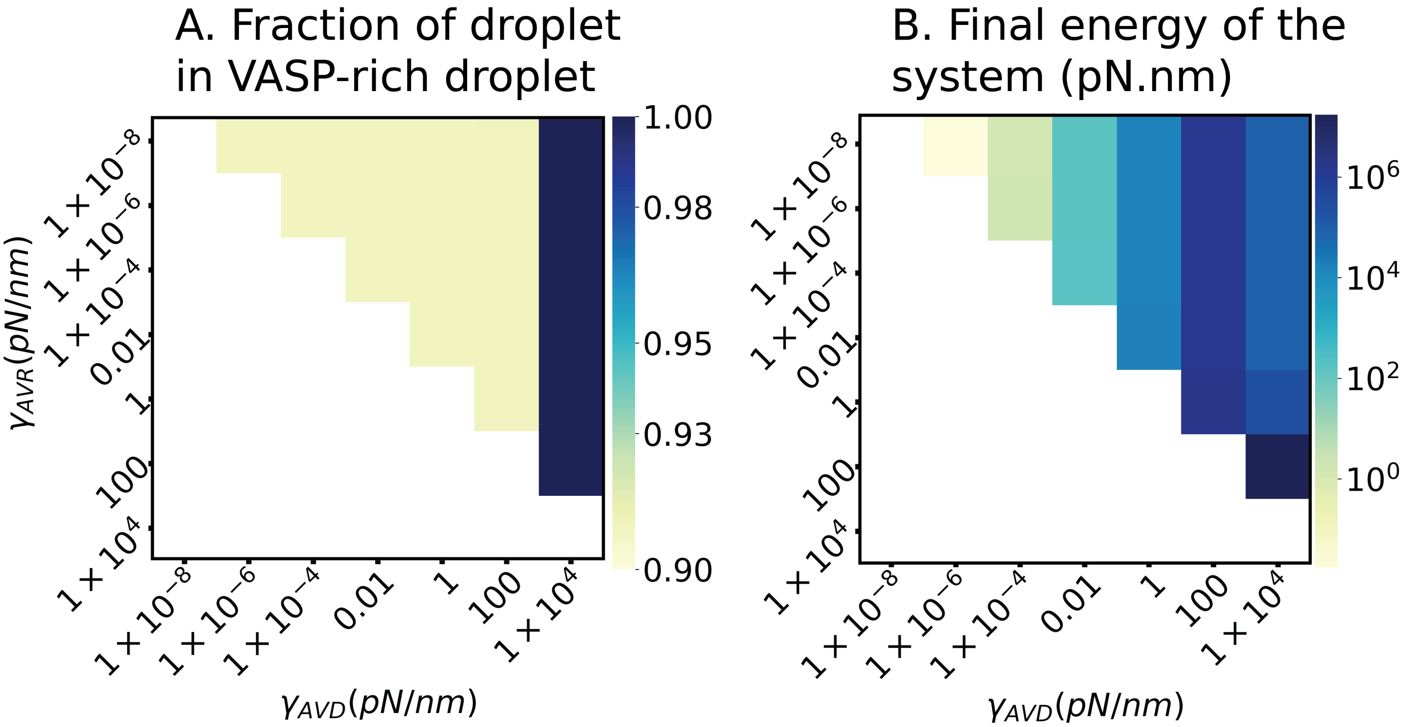


Supplemental Figure S 6: **Parameter sweep to determine the surface tension of the interface that bundled-actin ring shares with VASP-rich and VASP-dilute phases.** A) The fraction of actin bundle that is wetted by the VASP-rich droplet is shown. B) Heat map shows the energy of the system corresponding to the energy minimized configuration. As we are interested in the parameters that favor interaction of actin filaments with the VASP-rich phase, we only considered $\gamma_{AVR} < \gamma_{AVD}$

.

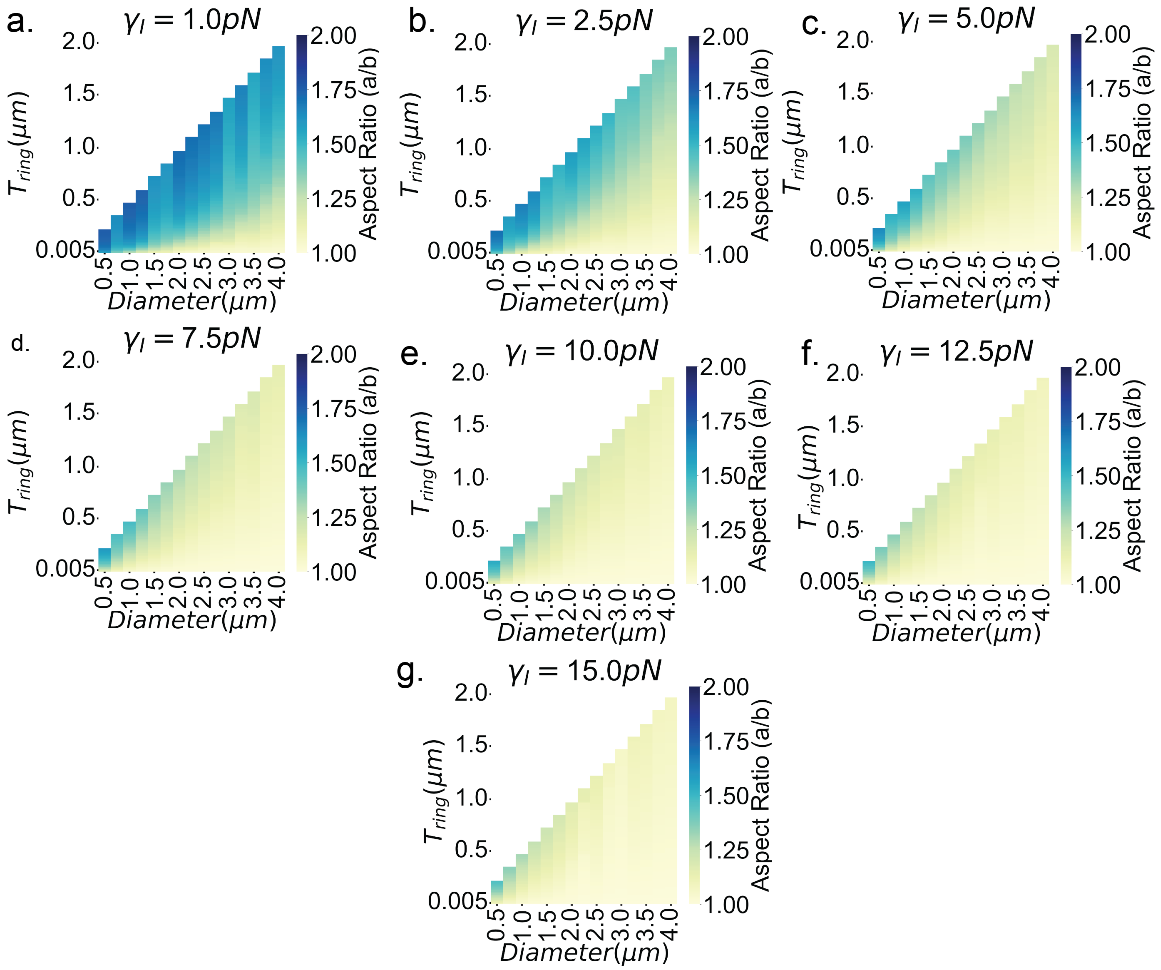


Supplemental Figure S 7: **Droplet shape depends on bundled-actin ring thickness and droplet shape change parameter,** $\gamma_I$ a-g. Each panel shows heat map of droplet aspect ratio at energy minimum as ring thickness of bundle is systematically varied (color bar shown to the right of each panel) at a given $\gamma_I$ value (mentioned on top). We find that lower $\gamma_I$ values of 1, and 2.5 $pN$ allow for larger aspect ratios of droplets while increasing the surface energy parameter leads to less-deformable droplets.

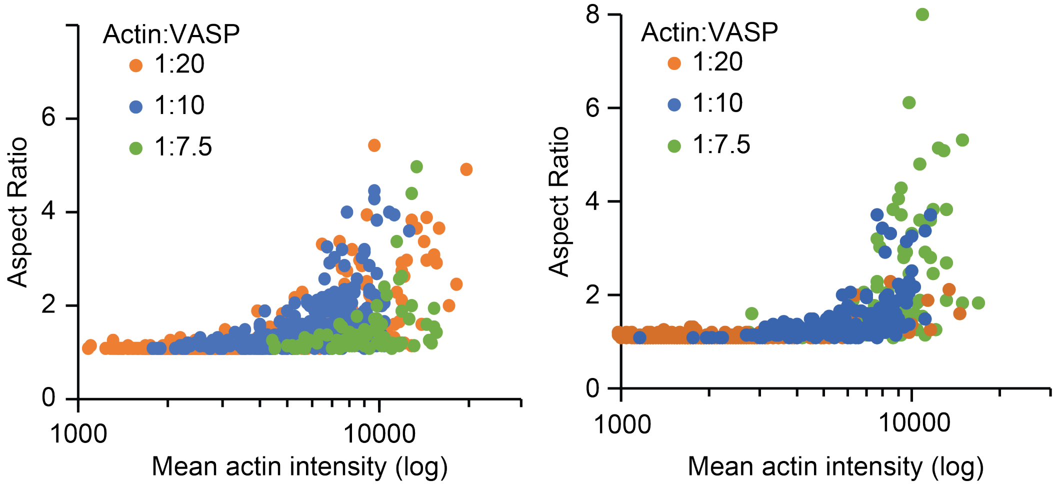


Supplemental Figure S 8: **Independent replicates of data shown in Figure 5b.** Left: n= 1196 droplets counted over at least 3 images per condition. Right: n = 594 droplets counted over at least 3 images per condition.

.

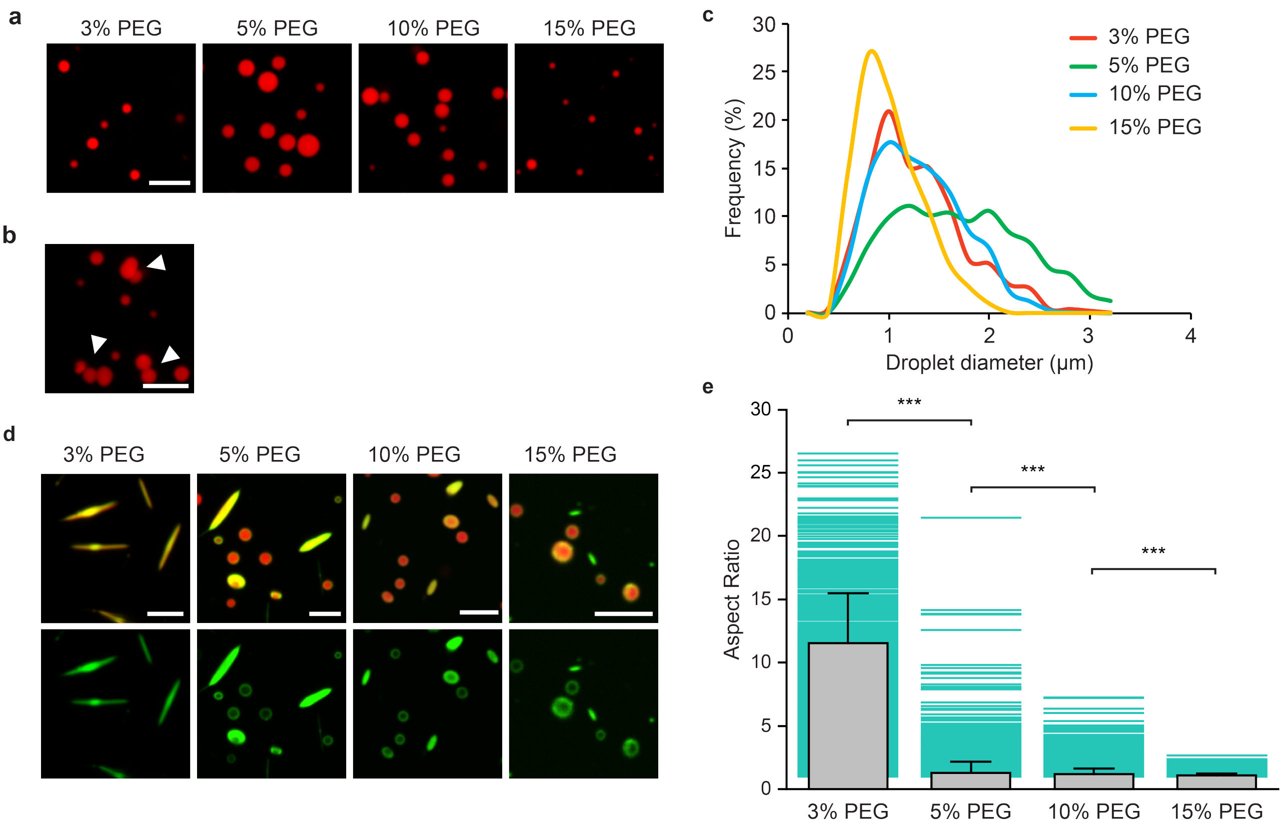


Supplemental Figure S 9: **A liquid-like state is required for droplet bundling.** (a) 10 $\mu M$ VASP droplets formed under increasing PEG. Scale bar 5 $\mu m$. (b) Droplets formed from 10% PEG are more solid than their 3% counterparts, and do not coalesce upon contact. Scale bar 5 $\mu m$. (c) Distribution of droplet size under increasing PEG concentrations. Data from n = 3 independent experiments with at least 3 images analyzed per condition. (d) Droplets formed from increasing PEG do not bundle actin into elongated structures. Scale bar 5 $\mu m$. (e) Distribution of droplet aspect ratios of 10 $\mu M$ VASP droplets containing 2 $\mu M$ actin as a function of PEG concentration. Bar data are mean $\pm$ SD, and teal background lines represent data points. For 3% PEG n = 877 droplets, for 5% PEG n = 2888 droplets, for 10% PEG n = 2802 droplets, for 15% PEG n = 2527 droplets across 3 independent experiments. Brackets indicate data that was tested for significance using an unpaired, two-tailed t-test. *** denotes $p < 0.001$.

#### 2 Supplementary Videos and Descriptions

**Supplemental Video S1.** VASP droplets fuse and re-round upon contact.

35 $\mu M$ VASP droplets (labeled with Alexa Fluor-647, shown in red) quickly fuse upon contact.

**Supplemental Video S2.** VASP droplets exhibit quick and complete fluorescence recovery after photobleaching.

VASP droplets (labeled with Atto-488, shown in red) were bleached at $t = 9s$. Fluorescence recovery occurred in approximately 5 minutes post-bleach.

**Supplemental Video S3.** Actin localized to VASP droplets distributes peripherally into rings over time.

2 uM actin (labeled with Atto-488, shown in green) added to 13 $\mu M$ VASP droplets is initially homogenous, but over time redistributes into a ring.

**Supplemental Video S4.** VASP droplets containing actin rings deform into rods.

15 $\mu M$ VASP (labeled with Atto-594, shown in red) droplets containing 3 $\mu M$ actin (labeled with Atto-488, shown in green). Actin is shown initially in a ring conformation and collapses into rods.

**Supplemental Video S5.** VASP droplets containing actin rings deform into rods that elongated into bundles.

10 $\mu M$ VASP (labeled with Atto-594, shown in red) droplets containing 2 $\mu M$ actin (labeled with Atto-488, shown in green). Actin is shown initially in a ring conformation and collapses into a rod that elongate bidirectionally into a bundle-like linear structure.

**Supplemental Video S6.** Elongation of VASP-actin bundles over time.

Bundles formed by 10 $\mu M$ VASP droplets (labeled with Alexa Fluor 647, shown in red) and 2 $\mu M$ actin (labeled with Atto-488, shown in green) elongate into linear structures over time.

**Supplemental Video S7.** Elongating droplet bundles zipper together upon contact.

Bundles formed by 10 $\mu M$ VASP droplets (labeled with Atto-647, shown in red) and 2 $\mu M$ actin (labeled with Atto-488, shown in green). Bundles elongate with time and upon contact, zipper together.

**Supplemental Video S8.** New actin monomers are added to the growing tip of the droplet bundles.

Bundles formed by 10 uM VASP droplets (labeled with Atto-594, shown in red) and 2 $\mu M$ actin (labeled with Atto-488, shown in green) were bleached at $t = 9s$. Increased fluorescence at the tips indicates new actin monomers are being added to the tip, while the polymerized actin along the shaft remains dark.

**Supplemental Video S9.** New actin monomers enter the transforming droplet over time.

An initially round droplet formed by 10 $\mu M$ VASP (labeled with AlexaFluor-647, shown in red) in the presence of 2 $\mu M$ actin (labeled with Atto-488, shown in green) was bleached at $t = 20s$. As the droplet transforms into an elongated bundle, both VASP and actin intensity increase over time, indicating that new proteins enter the droplet from the surrounding solution.

#### 3 Supplementary Methods - model development

When actin is added to VASP droplets, actin preferentially diffuses to the VASP-rich droplets where actin polymerization is enhanced due to VASP-actin interactions (Figure 2). As VASP-actin interactions are energetically favored, actin filaments never disassociate from the VASP-rich droplets. The preferred shape of the actin filaments that are constrained in a droplet of radius $R$ depends on the filament length, $L << L_p$ ($L_p$-persistence length). When $L < 2R$, filaments shapes are straight, subject to thermal fluctuations. As the filament grows, and reaches $L > 2R$, filaments assume the corresponding low energy configuration by following the curvature of the droplet forming a cortex-like shell (filaments with radii of curvature $< R$ have higher bending energy penalties). Finally, VASP-actin crosslinking-driven contractility brings the actin filaments together resulting in the formation ring-shaped actin bundles. Here, we model the shape changes resulting due to actin bundles that are thoroughly wet by VASP molecules in droplets by considering the mechanical free energies associated with both the actin bundle and the VASP droplet in two dimensions as shown in Figure S5a. Our model shows that as we increase the amount of actin in the ring, circular droplets readily deform to form elliptical droplets. Additionally, we see that the aspect ratio of droplets scales non-linearly with the bundled-actin ring thickness.

##### 3.1 Assumptions

We make the following assumptions in developing the model for a VASP droplet that contains an actin ring made of actin filaments that are bundled together by VASP.

- We model the droplet as a 2D circle; we assume that the bundled-actin ring is restricted to the plane of observation modeled here and does not undergo out-of-plane deformations.

- The surface area of the droplet is conserved for all values of ring thickness. This is consistent with experimental observations and stems from the underlying assumption that the number of VASP molecules is conserved given a droplet radius.

- The bundled-actin ring thickness is inelastic and does not change as a function of local stresses experienced by the ring. In other words, we assume that the filament-filament crosslinks that are part of the bundled-actin ring are extremely rigid. This assumption is justified, in part, as the actin ring exists in a droplet with excess VASP molecules.

- We assume that the bundled-actin ring undergoes complete wetting when exposed to either the VASP-rich and VASP-dilute phase.

- Bundled-actin ring thickness is a linear function of number of filaments and the persistence length of bundled actin ring scales linearly with the number of filaments in the ring.

##### 3.2 Table of Parameters

| Parameter | Value | Description | Reference |
|---|---|---|---|
| $D_{droplet}$ | $1-4\mu m$ | Droplet diameter | - |
| $2a, 2b$ | $4ab = D_{droplet}^2$ | Major and minor axis spans of ellipse obtained from deformation of circular droplet | - |
| $T_{ring}$ | Guided by experiments | The thickness of bundled-actin ring | - |
| $\delta$ | 15 nm | Spacing between actin filaments in a VASP crosslinked bundle ring (12nm spacing between actin binding domains in Ena) | [10] |
| $N_{fil}$ | $= \frac{T_{ring}}{\delta}$ | Number of filaments in the bundled -actin ring | - |
| $L_p$ | $= 17.73\mu m$ | Persistence length of single actrin filament | [11] |
| $L_{cyl}$ | $100nm$ | Length of cylindrical segment | - |
| $k_{stretch}$ | $N_{fil}.100pN/nm$ | Stretching energy constant of bundled -actin ring | [12] |
| $k_{bend}$ | $N_{fil}.\frac{L_p K_B T}{L_{cyl}} pN.nm$ | Bending energy of bundled-actin ring | Note1 [11] |
| $\gamma_{AVR}$ | $1 \times 10^{-8} pN/nm$ | Determined from simulations as described below. Wetting energy parameter for actin in VASP-rich phase | - |
| $\gamma_{AVD}$ | $1 \times 10^{4} pN/nm$ | Determined from simulations as described below. Wetting energy parameter for actin in VASP-dilute phase | - |
| $\gamma_I$ | Free parameter | Droplet surface energy parameter | - |

Table 1: Table of parameters used to simulate droplet shapes resulting from bundled-actin ring of varying thicknesses.

Note1: Bathe et al. showed that the bending stiffness $(k_b)$ depends on the bending stiffness of individual actin filaments $(k_{b,fil} = L_p.k_B T)$, number of filaments $(N)$, the inter-filament coupling parameter $(\alpha)$, crosslinker thickness parameter $(\chi)$, and the non-dimensional factor $(c(q_j))$ [13]. The expression is given by,

$$k_b = k_{b,fil} N \left( 1 + \frac{\chi^2 (N-1)}{1 + c(q_j) \frac{N + \sqrt{(N)}}{\alpha}} \right). \tag{S1}$$

As VASP-crosslinked actin networks produce weaker elastic responses than fascin and $\alpha$-actinin [14], we assume that the inter-filament coupling parameter is really weak. Hence, we assume that the contribution of nonlinear term in the above expression is low and hence consider the bending stiffness to scale linearly with number of filaments. The total energy of the bundled-actin ring interacting with a medium of VASP-rich and VASP-dilute regions is given by the sum of following free energies.

##### 3.3 Mechanical energy of the bundle

We begin by approximating the set of actin filaments found in the droplets as a single bundled ring as shown in Figure S5b. We choose to discretize the bundle into segments of length 100nm that can bend around hingepoints. The segment length is chosen to acoomodate for faithful reproduction of bundle curvatures at minimal computational overhead. Smaller segment lengths increase the computational overhead of the model while larger segment lengths suffer from poor approximations of bundle curvature. Consider $\mathbf{r_{bi}}$ to be the coordiante of bead $i$ in $D$ dimensions. Beads $\mathbf{r_{bi}}$ and $\mathbf{r_{bi+1}}$ constitute a bundle segment while the beads $\mathbf{r_{bi}}$, $\mathbf{r_{bi+1}}$ and $\mathbf{r_{bi+2}}$ constitute a hinge point. The mechanical energy of the bundle is given as,

$$E_{mech}(\mathbf{r}) \quad = \sum_{i,i+1} k_{stretch,i}\Bigg(\|\mathbf{r_{b,i}} - \mathbf{r_{b,i+1}}\|^2\Bigg) + \sum k_{bend,i,i+1,i+2} \frac{\big(\mathbf{r_{b,i}}-\mathbf{r_{b,i+1}}\big)\big(\mathbf{r_{b,i+1}}-\mathbf{r_{b,i+2}}\big)}{\|\mathbf{r_{b,i}}-\mathbf{r_{b,i+1}}\|\|\mathbf{r_{b,i+1}}-\mathbf{r_{b,i+2}}\|}. \tag{S2}$$

##### 3.4 Wetting energy of the bundled-actin ring

When the bundle interacts with the VASP-rich or with the VASP-dilute phase, the wetting free energy depends on the surface area of the bundle that is exposed to the solvent. If $T_{ring}$ is the radius of the bundle (or ring thickness) with a contour length, $L$, the wetting energy depends on the solvent that the bundle interacts with and is given by,

$$E_{wet} = \int_0^L T_{ring}\gamma(e)ds. \tag{S3}$$

As we assume total wetting of the bundled-actin ring by both VASP-rich and VASP-dilute phases, contact angle is 0. Please note that the surface tension is defined as a function of eccentricity of the droplet, $e$. To ensure a smooth transition of $\gamma$ around the interface, $\gamma(e)$ is defined as,

$$\Delta \quad = \frac{1}{2}\Bigg(\gamma_{AVR} - \gamma_{AVD}\Bigg), \tag{S4}$$

$$\gamma(e) \quad = \Delta\tanh(f(e)) + \Delta + \gamma_{AVD}. \tag{S5}$$

Figure 5c shows variation in surface tension across the interface for various values of f(e). In this study, we consider f(e)=64(e-1).

###### 3.4.1 Calculating $G_{wet}$

To numerically calculate $G_{wet}$ of a rod, we represent the equation of the rod connecting points $\mathbf{r_1}$ and $\mathbf{r_2}$ as,

$$\mathbf{r_p} = \mathbf{r_1} + s(\mathbf{r_2} - \mathbf{r_1}), 0 \leq s \leq 1. \tag{S6}$$

The equation of an ellipsoid can be written as,

$$(\mathbf{r} \odot \mathbf{r}).\mathbf{p} = 1. \tag{S7}$$

where $\mathbf{r} = x\hat{i} + y\hat{j}$ and $\mathbf{p} = a^{-2}\hat{i} + b^{-2}\hat{j}$ and $\odot$ represents the Hadamard product. In order to determine the wetting energy numerically, each rod is discretized into $N$ rod segments of length, $\delta L = L/N$. We calculate the eccentricity of each of the rod segments (average of the two ends) to numerically estimate the wetting energy of the rod.

###### 3.4.2 Choosing wetting energy parameters

We next determine the surface tension of the interface actin-ring creates with VASP-rich medium ($\gamma_{AVR}$) and the interface actin-ring creates with VASP-dilute medium ($\gamma_{AVD}$) by conducting a parameter sweep as these values are not readily experimentally measurable. First, an actin bundle of length $1.2\mu m$ was placed inside a cirular VASP droplet of diameter $1\mu m.$ The bundle was placed such that 100nm of actin is exposed to the VASP-dilute phase. Surface tension values were systematically tested to determine the values that lead to least energy penalty.

##### 3.5 Surface energy of the interface between VASP-rich VASP-dilute phases

While the total surface area of the droplet is held constant throughout our simulations, shape change happens through changes to the perimeter of the droplet as shown in Figure S5d. Consider an elliptical droplet whose parametric equation with $2a$ and $2b$ respectively as the major and minor axis span is given by,

$$x = acos(\theta), y = bsin(\theta). \quad \text{(S8)}$$

(S9)

The perimeter of an ellipse, $L$ is given by,

$$L = 4\int_0^{\pi/2} \sqrt{((x'(\theta))^2 + (y'(\theta))^2)} d\theta. \quad \text{(S10)}$$

The perimeter is given by an infinite series as follows

$$L = \pi(a+b)\sum_{n=0}^{\infty} \binom{0.5}{n}^2 h^n, \quad \text{(S11)}$$

where $h = (\frac{a-b}{a+b})^2$. In this study, we consider the first seven terms in the expansion. The surface free energy of the droplet with perimeter L is given as,

$$\Delta E_{surface} = \gamma_I L, \quad \text{(S12)}$$

where $\gamma_I$ represents the surface tension of the interface between the VASP-rich and VASP-dilute phases.

##### 3.6 Simulation protocol

The total mechanical free energy of bundled-actin ring in phase separated VASP system is computed based on the interactions described above. We investigated how the aspect ratio of VASP droplets changes at various actin bundle thicknesses. We assume that the bundled-actin rings are flat, isotropic sheets of actin. As the G-actin binding domains of VASP monomers are separated by $\approx 12\ nm$ [15], we assume an average spacing of 15 nm between actin filaments in the ring. As a result, given a ring thickness, we can estimate the number of filaments in the ring and assume a linear relationship between number of filaments ($N_{fil}$) and bending stiffness of the ring $K_{b,ring} = K_{b,fil} \times N_{fil}$. The role of nonlinear relationship between bundle stiffness and number of filaments will be explored is an important future direction to pursue.

To understand how ring thickness affects shapes of droplets, we begin by generating an actin ring made of single actin filament through an iterative extension-minimization protocol described below. We begin with a VASP-rich droplet of diameter D surrounding an actin filament of length $L = D$. We then generate an actin ring by iteratively minimizing the total energy of the system by extending the actin filament length by $0.1D$ each time for 21 cycles $L_{final} = 3.1D \approx \pi D$. The total energy of the system is minimized using a nonlinear optimization algorithm called Sequential Least Squares Programming (SLSQP) with a tolerance value of $10^{-6}$ under constant droplet area constraint. Droplet shapes with bundled-actin rings of varying thickness are then considered by iteratively increasing the ring thickness through $N_{fil}$ and $k_{b,ring}(N_{fil})$ to determine the minimum energy configuration at each step. Between each minimization, random noise is added (uniform distribution between 0 and 0.1 nm) to the coordinates to ensure robust minimization. This perturbation procedure also enables us to study multiple replicates.

##### 3.7 Derivation of the scaling relationship showing the maximum bundle thickness above which a circular droplet deforms into an ellipse

We build on the scaling relationship derived by Limozin et al. [16]. Limozin et al. studied the shape of actin bundles deposited on the inner surfaces of lipid vesicles. We adapt the same arguments for liquid droplets. Consider a droplet of radius R with an actin bundle of length $L$. As the bundle is subject to thermal fluctuations in the droplet, we approximate the shape of the bundle as fluctuations around the droplet radius, $R = D_{droplet}/2$. Expressing the radial coordinate as a function of arc length $s$,

$$r(s) = R + u(s), u(\theta) = \sum_m u_m cos(m\theta), s = R\theta. \quad \text{(S13)}$$

To understand how the bundle thickness is related to droplet radius, authors derive the bending free energy of the bundle and use equipartition theorem to obtain the scaling relationship.

##### 3.8 Derivation for the bending free energy of the bundle

In this section, we re-derive the scaling expression shown by Limozin et al. The bending free energy of a bundle with local curvature $\kappa(s)$, and flexural rigidity $k_b$ is given by,

$$E_{bend} = \frac{k_b}{2}\int_0^L \kappa(s)^2 ds. \tag{S14}$$

Using the derivation in Section 3.10, we assume that the bundle traces the curvature of the elliptical droplet and write the curvature of the bundled-actin ring as,

$$\begin{aligned}
\kappa(s)^2 &= \Bigg(\Big(r^{''} - \tfrac{r}{R^2}\Big)^2 + (\tfrac{2r^{'}}{R})^2\Bigg),\\
r^{'}(s) &= \tfrac{1}{R}\sum m u_m sin(\tfrac{ms}{R}),\\
r^{''}(s) &= \tfrac{-1}{R^2}\sum m^2 u_m cos(\tfrac{ms}{R}),
\end{aligned}$$

$$r^{'}(s)^2 = \frac{1}{R^2}\Bigg(\sum_m m u_m sin(\frac{ms}{R})\Bigg)^2. \tag{S15}$$

Assuming that on an average, the different modes $m$ are uncorrelated, we can ignore the cross-correlation between different modes. Hence, we write the average value as,

$$\langle r^{'}(s)^2\rangle = \frac{1}{R^2}\sum_m \Bigg\langle m^2 u_m^2 sin^2(\frac{ms}{R})\Bigg\rangle. \tag{S16}$$

Using a similar expression for $r^{''}(s)$, we write the average squared curvature of the bundle as,

$$\langle\kappa(s)^2\rangle = \frac{1}{R^2} + \frac{2}{R^3}\sum_m u_m\langle cos(\frac{ms}{R})(1+m^2)\rangle + \frac{1}{R^4}\sum_m\Bigg((1+m^4)u_m^2\langle cos^2(\frac{ms}{R})\rangle + 2u_m^2\langle sin^2(\frac{ms}{R})\rangle\Bigg). \tag{S17}$$

Please note that the curvature value is primarily determined by the shape of the droplet ($1/R^2$). Subsequent corrections to the curvature are given by terms with $1/R^3$ and $1/R^4$. We investigate the contribution of each of these terms to the bending free energy of the bundle. The bending free energy of the actin bundle is given by,

$$E_{bend} = \frac{k_b}{2}\int_0^L \langle\kappa(s)^2\rangle ds. \tag{S18}$$

Substituting expression for curvature in Eq. S17, we find three key integrals as shown below.

$$\begin{aligned}
\int_0^L cos^2(\frac{ms}{R})ds &= \textstyle\int_0^L 1 + cos(\frac{2ms}{R})ds = L - \frac{R}{2m}sin(\frac{2mL}{R}),\\
\int_0^L sin^2(\frac{ms}{R})ds &= \textstyle\int_0^L 1 - cos(\frac{2ms}{R})ds = L + \frac{R}{2m}sin(\frac{2mL}{R}),
\end{aligned} \tag{S19}$$

and

$$\int_0^L cos(\frac{ms}{R})ds = -\frac{R}{m}sin(\frac{ms}{R}). \tag{S20}$$

Substituting these terms in S17 and also considering that the fluctuations from sin and cos terms will dampen on average, we get,

$$E_{bend} = \frac{kb}{2}\Bigg(\frac{L}{R^2} + \frac{L}{R^4}\textstyle\sum_m (2+m^4)u_m^2\Bigg). \tag{S21}$$

Equation S21 shows that the primary contribution to bending energy of the bundle is from $L/R^2$ due to the confinement of bundle inside the droplet. Ignoring the bending free energy from droplet confinement and consider the excess bending free energy, denoted as $\Delta E_{bend}$,

$$\Delta E_{bend} = \frac{k_b}{2}\frac{L}{R^4}u_m^2\big(2 + m^4\big). \tag{S22}$$

##### 3.9 Scaling relationship

Equipartition theorem suggests that each mode contributes $\frac{k_B T}{2}$ energy, where $k_B$ is the Boltzmann constant and $T$ is the temperature. Thus, each term within the summation contributes $\frac{k_B T}{2}$. Ignoring the mode-dependant contribution $f(m)$, we get that,

$$\frac{k_B T}{2} = \frac{k_b}{2} f(m)\frac{L}{R^4}u_m^2. \tag{S23}$$

Assuming that the thickness of the cortex is proportional to the Fourier amplitude, we get,

$$u_m \approx \frac{R^2}{\sqrt{(LL_p)}}. \tag{S24}$$

where $L_p = k_b/k_B T$ is the persistence length of the bundle with flexural rigidity, $k_b$. Considering actin segment of length $L = 2\pi R$, the scaling relationship is given by,

$$u_m \approx \frac{R^{1.5}}{\sqrt{(L_p)}}. \tag{S25}$$

##### 3.10 Derivation for the curvature, $\kappa(s)$

Let us denote the coordinates of the actin bundle, and the arc length, s are given as,

$$x = rcos(\theta), y = rsin(\theta), s = R\theta,$$
$$x(s) = r(s)cos(\frac{s}{R}), y(s) = r(s)sin(\frac{s}{R}).$$

Differentiating twice with respect to s,

$$\begin{aligned}
x^{'}(s) &= r^{'}cos(\tfrac{s}{R}) - \tfrac{r}{R}sin(\tfrac{s}{R}),\\
y^{'}(s) &= r^{'}sin(\tfrac{s}{R}) + \tfrac{r}{R}cos(\tfrac{s}{R}),\\
x^{''}(s) &= r^{''}cos(\tfrac{s}{R}) - 2\tfrac{r^{'}}{R}sin(\tfrac{s}{R}) - \tfrac{r}{R^2}cos(\tfrac{s}{R}),\\
y^{''}(s) &= r^{''}sin(\tfrac{s}{R}) + 2\tfrac{r^{'}}{R}cos(\tfrac{s}{R}) - \tfrac{r}{R^2}sin(\tfrac{s}{R}).
\end{aligned}$$

The curvature, $\kappa(s)$ is given by,

$$\begin{aligned}
\kappa(s) &= \sqrt{\Big((x^{''}(s))^2 + (y^{''}(s))^2\Big)},\\
\kappa(s)^2 &= \Big(\big(r^{''} - \tfrac{r}{R^2}\big)^2 + (\tfrac{2r^{'}}{R})^2\Big)\Big(cos^2(\tfrac{s}{R}) + sin^2(\tfrac{s}{R})\Big).
\end{aligned}$$

##### 3.11 Exploring the role of bundle bending and droplet surface energies in ring-rod transition

Experimental evidence shows transition of 2D rings in circular droplets to 1D bundles embedded in an elongated ellipsoidal droplets leading to higher aspect ratios. (Figures 4a, and 5a-d) As this transition is mediated by bending energy of the bundle and the surface energy of the droplet, we chose to understand these as a function of aspect ratio. To understand the energetic contribution of relevant energies as the aspect ratio increases, we employ a continuum model of 2D droplet with a bundle that traces the curvature of the ellipse. The bending energy of the bundle of length L in a droplet of radius $R_{droplet} = 1\mu m$ is given by $E_{bend} = k_b/2 \int_0^L \kappa(s)^2 ds$, where $\kappa(s)$ is the local curvature of the actin bundle with flexural rigidity $k_b$ at a given arc length, $s$. The surface energy of the VASP-droplet with a surface energy parameter, $\gamma_I$ is given by $E_{surf} = \gamma_I P$, where $P$ is the perimeter of the elliptical droplet. We calculate the energetic landscape as the aspect ratio of the droplet is increased from 1 (circle) to the maximum value $\pi^2$ (rod) under constant area constraint, *i.e.* $ab = R^2_{droplet}$. For sake of simplicity, we assume $k_b = 1$ and $\gamma_I = 1$. In Figure 5c, we show that as the aspect ratio of a droplet is increased, droplet deforms from a circle to an ellipse with a larger perimeter leading to a steady increase in the surface energy. Additionally, we see that the bending energy of the droplet increases steadily initially and the bending energy reduces at high aspect ratios $(> 9)$. Figure 5d shows configurations of bundles at various aspect ratios. We see that the initial increase in bending energy stems primarily from bundle segments found at the ends of the droplet major axis. As the aspect ratio is increased, the bundle energy continues to increase till $P(a/b) < 2L$. When $P(a/b) > 2L$, bundle entirely exists in the lower curvature regions of the droplet leading to decrease in bundle bending energy. This suggests a plausible mechanism for the unwinding of a circular ring into a linear rod while being constrained to a plane within the droplet. The role of droplet dimensionality and actin bending rigidity need further exploration.
